# Large-scale transcriptomics to dissect two years of the life of a fungal phytopathogen interacting with its host plant

**DOI:** 10.1101/2020.10.13.331520

**Authors:** Elise J. Gay, Jessica L. Soyer, Nicolas Lapalu, Juliette Linglin, Isabelle Fudal, Corinne Da Silva, Patrick Wincker, Jean-Marc Aury, Corinne Cruaud, Anne Levrel, Jocelyne Lemoine, Regine Delourme, Thierry Rouxel, Marie-Hélène Balesdent

**Affiliations:** Université Paris-Saclay, INRAE, AgroParisTech, UMR BIOGER, 78850 Thiverval-Grignon, France; Génomique Métabolique, Genoscope, Institut François Jacob, CEA, CNRS, Université d’Evry, Université Paris-Saclay, 91057 Evry, France; Genoscope, Institut François Jacob, CEA, Université Paris-Saclay, Evry, France; INRAE, Institut Agro, Univ Rennes, IGEPP, 35653, Le Rheu, France

## Abstract

The fungus *Leptosphaeria maculans* has an exceptionally long and complex relationship with its host plant, *Brassica napus*, during which it switches between different lifestyles, including asymptomatic, biotrophic, necrotrophic, and saprotrophic stages. The fungus is also exemplary of “two-speed” genome organisms in which gene-rich and repeat-rich regions alternate. Except for a few stages of plant infection under controlled conditions, nothing is known about the genes mobilized by the fungus throughout its life cycle, which may last several years in the field. We show here that about 9% of the genes of this fungus are highly expressed during its interactions with its host plant. These genes are distributed into eight well-defined expression clusters, corresponding to specific infection lifestyles or to tissue-specific genes. All expression clusters are enriched in effector genes, and one cluster is specific to the saprophytic lifestyle on plant residues. One cluster, including genes known to be involved in the first phase of asymptomatic fungal growth in leaves, is re-used at each asymptomatic growth stage, regardless of the type of organ infected. The expression of the genes of this cluster is repeatedly turned on and off during infection. Whatever their expression profile, the genes of these clusters are located in regions enriched in heterochromatin, either constitutive or facultative. These findings provide support for the hypothesis that fungal genes involved in niche adaptation are located in heterochromatic regions of the genome, conferring an extreme plasticity of expression. This work opens up new avenues for plant disease control, by identifying stage-specific effectors that could be used as targets for the identification of novel durable disease resistance genes, or for the in-depth analysis of chromatin remodeling during plant infection, which could be manipulated to interfere with the global expression of effector genes at crucial stages of plant infection.

**Author Summary:** Fungi are extremely important organisms in the global ecosystem. Some are damaging plant pathogens that threaten global food security. A knowledge of their biology and pathogenic cycle is vital for the design of environmentally-friendly control strategies. Unfortunately, many parts of their life cycle remain unknown, due to the complexity of their life-cycles and technical limitations. Here, we use a rapeseed pathogen, *Leptosphaeria maculans*, which has a particularly complex life-cycle, to show that large-scale RNA-Seq analyses of fungal gene expression can decipher all stages of the fungal cycle over two years of interaction with living or dead hosts, in laboratory and agricultural conditions. We found that the fungus uses about 9% of the genes of its genome specifically during interactions with the plant, and observed waves of extremely tight, complex regulation during the colonization of specific tissues and specific parts of the life-cycle. Our findings highlight the importance of genes encoding effectors, small secreted proteins manipulating the host. This work opens up new avenues for plant disease control through the identification of stage-specific effectors leading to the discovery of novel durable disease resistance genes, or the analysis of epigenetic regulation, which could be manipulated to interfere with effector gene expression.

## Introduction

Fungi play a vital role in global ecosystems. They have highly complex life-cycles that often remain poorly understood, due to the multiplicity of fungal species and their behavior. Plant-associated fungi (endophytes, symbionts and phytopathogens) contribute to this complexity and diversity of behavior, as they may have beneficial, neutral or detrimental interactions with their hosts; some are of paramount importance in favoring the growth of plants or their adaptation to the environment, whereas others are highly damaging pathogens of cultivated plants [1].

Many fungal pathogens of annual crops display contrasting infection strategies (e.g. the biotrophs *Blumeria graminis* and *Ustilago maydis,* the hemibiotroph *Colletotrichum higginsianum* or the necrotroph *Botrytis cinerea*), with reproducible short cycles of colonization/sporulation, lasting from hours to weeks, on plant leaves. For such species, high-throughput RNA sequencing has made it possible to describe the stage-specific expression of genes during the few days or weeks over which the infection process occurs, under laboratory conditions. Such studies have identified concerted waves of pathogenicity gene expression during fungal infection [2–3], sets of genes involved in the switch from asymptomatic to necrotrophic growth in the hemibiotrophic fungus *Zymoseptoria tritici* [4,5], or four waves of expression for genes encoding putative effectors in *C. higginsianum* [6]. Short life-cycles, of no more than a few weeks on the host, are compatible with the description of biological and phytopathogenic features in experiments performed in controlled conditions.

In many other models, including symbionts and endophytes, the fungi spend much longer interacting with their host, and make use of different pathogenic/symbiotic developmental programs to adapt to different lifestyles during host colonization (this is the case in *Leptosphaeria maculans*, the fungus studied here), different host tissues (e.g. *U. maydis*; [7]), or different plant species (e.g. rust species with complex pathogenic cycles including two alternate host species; [8]). For these species, RNA-Seq approaches also seem to be the approach of choice for dissecting the entire fungal life-cycle and describing the biology of the fungus, together with the sets of genes involved and their regulation. However, the use of RNA-Seq to describe the fungal life-cycle is challenging and technically demanding. Even for fungal species that can be grown in axenic media and for which miniaturized pathogenicity assays have been developed, parts of the pathogenicity cycle remain inaccessible and cannot be reproduced in laboratory conditions. It is, therefore, essential to sample and monitor isolates from the wild. The second challenge is developing a sufficiently comprehensive *a priori* knowledge of all the relevant stages of the fungal life-cycle, and a means of sampling all of these stages for analysis. The third challenge is the generation of sufficient amounts of high-quality samples (often corresponding to minute amounts of fungal organisms in large amounts of plant material, or samples obtained blind, without knowing whether the fungus is present in the plant at the time of sampling) for RNA-Seq, to ensure the generation of enough reads for appropriate statistical analyses. The fourth challenge is coping with complex field samples, containing not only the phytopathogen and the host plant, but also many other fungal or bacterial species, necessitating a careful evaluation of the specificity of RNA-Seq mapping before analysis. The final challenge is coping with field samples subject to natural infection in variable environmental conditions, potentially resulting in a high level of heterogeneity in terms of the presence of the fungus and its developmental stage within the tissues, and attaining a biologically relevant continuity when monthly samples are obtained over a whole growing season.

*L. maculans*, a pathogen of rapeseed (*Brassica napus*), has an exceptionally complex life cycle compared to most other fungal phytopathogens attacking annual plants. It can infect different plant tissues, undergo multiple switches from biotrophy to necrotrophy during a lengthy lifespan in plant tissues, and can also live as a saprophyte on plant residues. The epidemiological cycle of *L. maculans* is well described [9,10]. It begins with the hemibiotrophic colonization of young leaves by spores generated by sexual reproduction (ascospores) in early autumn (October-November) in Europe. The fungus first colonizes the leaf tissues as a biotroph, for a few days or weeks, depending on the climatic conditions, without causing any symptoms. It then induces the development of necrotrophic leaf lesions in which its asexual spores (conidia) are produced. The fungus then migrates, without causing symptoms, from the petiole to the stem, where it lives in the plant tissues, as an endophyte, for several months. Finally, at the end of the growing season (May to July in Europe), it switches back to necrotrophic behavior, inducing the formation of a damaging stem canker that may result in plant lodging. Having completed all these stages of infection on living plant tissues, *L. maculans* then switches to a saprotrophic lifestyle, living on crop residues for up to three years. It develops structures for sexual reproduction to create the new inoculum (ascospores) for subsequent seasons on these residues. Due to its length and complexity, only a limited part of the life-cycle of *L. maculans* is amenable to laboratory experiments. Most of the RNA-seq-based time-course experiments published to date were performed during the first two weeks of infection, on cotyledons in controlled conditions [11–13], or in one set of stem infection conditions in controlled conditions [14].

In many models analyzing primary leaf infection, RNA-Seq approaches have highlighted the importance of genes encoding small secreted proteins (SSPs), acting as effectors, which often compromise plant defense responses [15]. In *L. maculans*, SSP genes are strongly expressed during cotyledon infection in controlled conditions [11–13], whereas another set of putative effectors and a toxin encoded by a secondary metabolite gene cluster, sirodesmin PL, are recruited during stem infection [14,16]. To date, 20 effector genes have been characterized in *L. maculans*, corresponding to the most documented class of pathogenicity genes in this species. These genes include nine cloned “early” effector genes (*AvrLm* genes) [17–25] located in AT-rich regions of the fungal genome enriched in transposable elements (TEs) [26] and encoding typical SSPs with few, if any, matches to sequences in databases, frequently enriched in cysteine residues. These genes are expressed and act during asymptomatic cotyledon infection. Eleven “late” putative effector genes, with molecular features similar to those of “early” effectors, were then shown to be expressed during stem colonization [14]. Unlike “early” effectors, these genes are located in gene-rich regions of the genome.

In addition to effector genes, the possible role in virulence of dozens of genes of different functional categories has been studied in *L. maculans*. These genes encode proteins involved in the biosynthesis of two secondary metabolites: abscisic acid (ABA) [27] and the toxin sirodesmin [16], and genes disrupted or silenced in *L. maculans,* 11 of which have been implicated in fungal virulence [28–36]. Again, all these studies were performed in controlled conditions, generally with miniaturized pathogenicity tests on cotyledons, and nothing is known about the involvement of the candidate genes, or of other pathogenicity genes, in other parts of the fungal life cycle. ‘Late’ effectors have been identified only in controlled conditions, and their involvement in the lengthy process of stem colonization in the field requires further investigation.

*L. maculans* has a well-defined bipartite genome consisting of gene-rich, GC-equilibrated regions and large AT-rich regions enriched in transposable elements (TE) but depleted of genes. The AT-rich regions account for 34.8% of the genome and are enriched in H3K9me3 (trimethylation of the lysine 9 residue of histone H3) marks, typical of constitutive heterochromatin in other species, whereas the GC-rich regions are enriched in H3K4me2 (dimethylation of the lysine 4 residue of histone H3) marks, typical of euchromatin [37–38]. In addition, species-specific genes and genes silenced during axenic growth display an enrichment in H3K9me3 and H3K27me3 (trimethylation of the lysine 27 residue of histone H3; [38]) heterochromatin modifications. The chromatin-based regulation of the expression of ‘early’ effector genes has been experimentally validated for a few *AvrLm* genes in this model [37].

We hypothesized here that, with the use of careful experimental and sampling procedures, it should be possible to use a large-scale transcriptomic approach to decipher the whole of the complex life cycle of *L. maculans* in interaction with its host, in both controlled and field conditions, and over time scales ranging from weeks (controlled condition experiments) to months or years (follow-up of plant infection on leaves and stems for a whole growing season, survival of *L. maculans* on residues remaining on the soil for one year). Our objective was, thus, to use RNA-Seq data to describe the complete fungal life-cycle and pathogenicity cycle and to identify the genes and functions involved in each stage. We found that about 9% of the genes of the fungus were expressed specifically during host-fungus interactions. A clustering of all the genes involved in the host-fungus interaction led to the detection of eight major waves of gene expression that could be consistently related to the consecutive stages of the *L. maculans* life-cycle, to tissue-specific expression, or corresponded to genes recycled for all similar lifestyles at different stages of the fungal life-cycle. The regulation of expression was strongly associated with the hosting of genes of the eight waves in heterochromatin landscapes, and heterochromatin dynamics is thought to be instrumental in allowing the recycling of expression for genes from the same wave during all stages of fungal biotrophy.

## Results

### Biological and RNA samples representing all aspects of the life-cycle of the fungus in interaction with its host

The RNA-Seq experiment was designed to generate samples corresponding to all stages of the interaction of *L. maculans* with its host plant, either alive (**Fig 1E-C**) or dead (stem residues after harvest; **Fig 1D**), in controlled conditions or in field experiments under natural inoculum pressure, over periods of time ranging from a few days to months or years (**Fig 1**; For a detailed description of biological samples, see the M&M section). As a reference, we included a series of samples grown in axenic conditions promoting a particular fungal developmental stage (mycelium growth, pycnidium differentiation, pseudothecium differentiation, resting conidia, conidial germination) (**Fig 1E**), to distinguish between these fundamental biological processes and those specific to interaction with the plant. In total, 102 biological samples, corresponding to 37 different sets of conditions, were subjected to RNA-Seq. Several quality control checks and parameter optimization were required before this heterogeneous dataset could be subjected to statistical analyses. *In fine,* 32 sets of conditions were retained for statistical analyses, whereas five sets of conditions were excluded because too few fungal reads were available (**S1 Text**, **S1** and **S2 Tables**).

**Fig 1.**
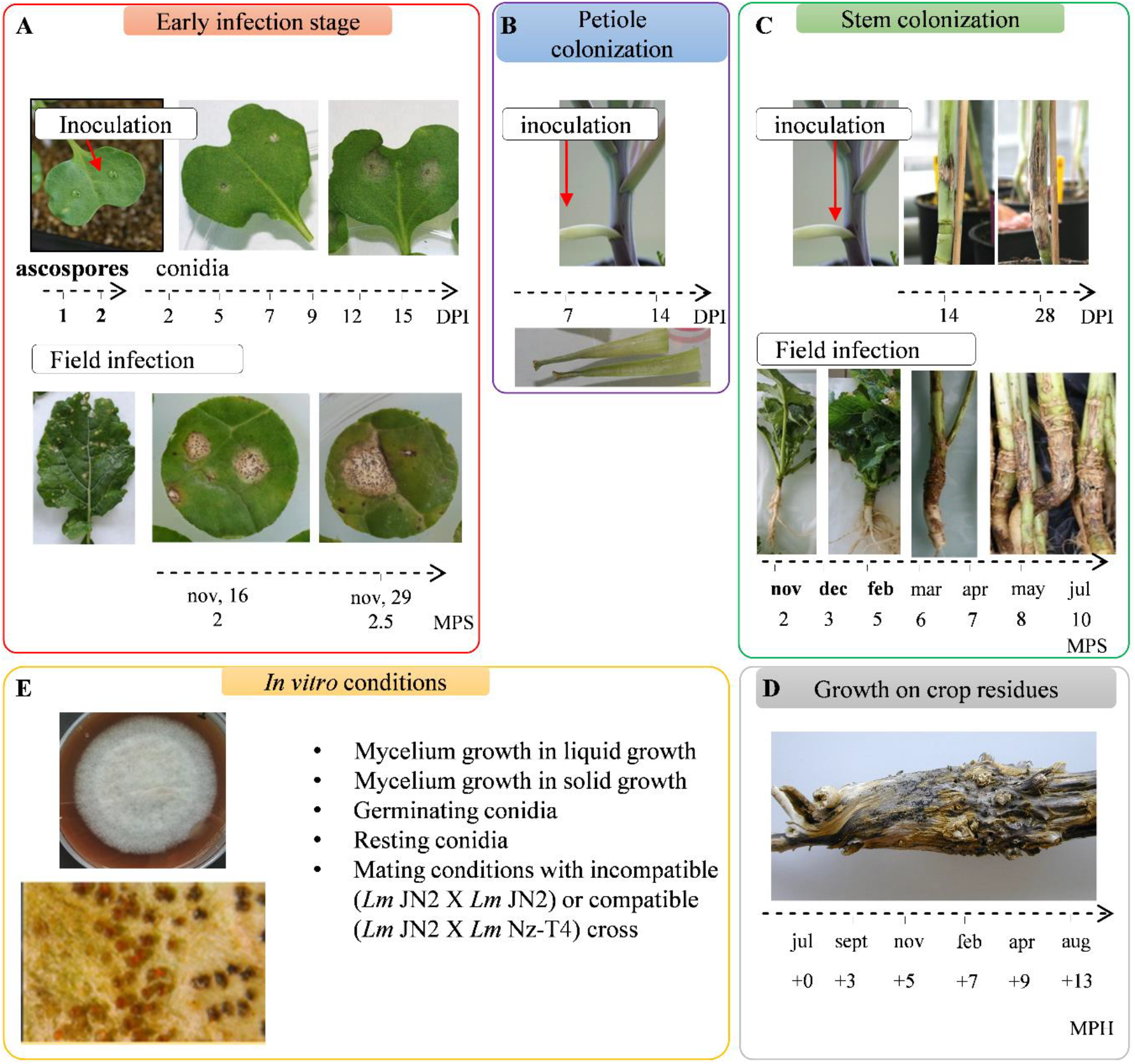
Conditions and samples used for the RNA-Seq study of the *Leptosphaeria maculans* life cycle. Time-course studies are represented by an arrow and the sampling time points are indicated. (A) RNA was extracted from cotyledons infected with ascospores one or two days post inoculation (DPI). Another time-course study was conducted by inoculation with conidia, with RNA extraction at six time points, from 2 to 15 DPI. Young leaves from naturally infected fields were sampled 2 or 2.5 months post sowing (MPS). (B) Petiole colonization was achieved on petioles inoculated in controlled conditions, with RNA extraction at 7 and 14 DPI. (C) Stem colonization samples were obtained in controlled conditions, at 14 and 28 DPI. Stem base samples from naturally infected fields were collected and used for RNA extraction every two to three months over an entire growing season. (D) Leftover stem residues were sampled every two to three months for one year after harvest (dates expressed in months post harvest, MPH). (E) We used 10 sets of *in vitro* growth conditions, based on two different media (liquid or solid) and corresponding to different physiological states (mycelium, sporulation, germinating and resting conidia, *in vitro* crossing conditions). The samples highlighted in bold did not provide enough fungal RNA-Seq reads for statistical analysis.

### Quality control analyses highlight the consistency of the RNA-Seq dataset

Clustering based on pairwise Pearson’s correlation analyses of the 32 sets of conditions identified four groups of clustered conditions (**Fig 2**). In each of these four groups, principal component analysis (PCA) showed a low level of variation between biological replicates, except for a few field samples and *in vitro* crosses (**S2 Text**, **S1A-D Fig**). The four clusters (**Fig 2**) corresponded to: (i) the early infection stage on cotyledons and leaves and colonization of the petiole (group 2), (ii) the late infection stage on stems (group 1), (iii) the saprophytic lifestyle (group 3), in controlled conditions or in the field, and (iv) all the *in vitro* growth conditions promoting the differentiation of pseudothecia and/or pycnidia (group 4).

**Fig 2.**
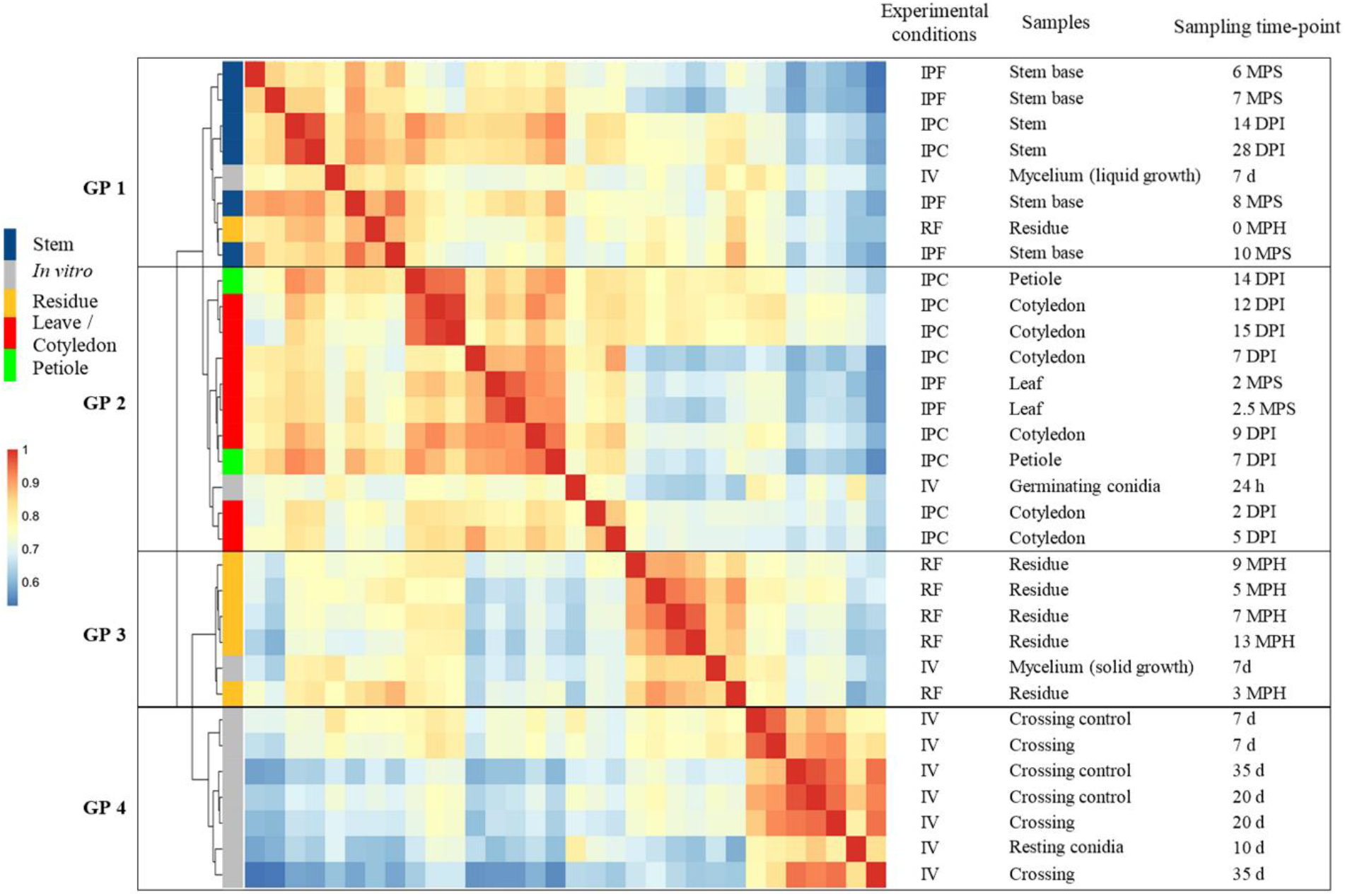
Hierarchical clustering of *Leptosphaeria maculans* gene expression in 32 conditions representing the fungal life cycle. Each gene with an FPKM count > 2 in at least one set of conditions was retained for the correlation analysis. A pairwise Pearson’s correlation analysis was performed between the 32 sets of conditions on Log_2_(FPKM+1) values. The resulting Pearson’s correlation matrix was subjected to hierarchical clustering (represented by the tree) with the ward.D2 method. The Pearson’s correlation results are shown on a color scale. Four groups of clustered samples were identified (GP1 to GP4). The experimental conditions are indicated as IPF, *in planta* field conditions; IPC, *in planta* controlled conditions; IV, *in vitro* conditions and RF, residues in field conditions. The infected plant tissues collected or the conditions under which the *in vitro* samples were obtained are detailed, together with the corresponding time points for sampling (DPI, days post-inoculation; MPS, months post-sowing; MPH, months post-harvest; d, days; h, hours).

Within each group, lifestyle provided a second level of sample discrimination, with samples from biotrophic or endophytic stages distinguished from necrotrophic-stage samples (**Fig 2**). For example, within group 2, samples displaying necrotrophic behavior on petioles 14 days post infection (DPI) were more closely related to the necrotrophic samples collected from cotyledons 12 and 15 DPI than to the corresponding biotrophic samples (petioles 7 DPI) (**Fig 2**). Similarly, in group 1, the residues collected immediately after harvest grouped with the last samples collected from the stem base, even though the residues did not originate from the same growing season, or from the same rapeseed variety, suggesting that the transition from stem necrosis to saprophytic life takes place after harvest (**Fig 2**).

Some axenic growth conditions were also grouped with some *in planta* conditions. Within group 2, germinating conidia grouped with the earliest samples collected from cotyledons, consistent with the use of conidia to inoculate plant tissues in controlled conditions (**Fig 2**). Samples grown in axenic culture on solid V8 medium under conditions promoting mycelial growth over sporulation clustered within group 3, indicating that mycelial growth is a major component of the saprophytic lifestyle in the wild. By contrast, mycelium grown in static liquid medium grouped with all conditions in which the fungus colonized stem tissues, possibly reflecting growth under low-oxygen conditions in plant vessels or intercellular spaces.

### Analytical strategy for identifying genes involved in pathogen-plant interactions

For the identification of fungal genes specifically involved in at least one stage of the pathogen-plant interaction, we used the 10 sets of *in vitro* growth conditions as 10 sets of control conditions, to exclude all genes involved in basic fungal metabolism or life traits other than those associated with plant infection (**Fig 3**). Using very stringent criteria (LogFC > 4; *p* value < 0.01), we focused on genes specifically overexpressed *in planta* relative to *in vitro* control conditions. We analyzed the expression patterns of these genes, their functional annotation, their genomic localization, and their location with respect to histone modifications during axenic growth. We then focused on genes encoding SSPs from the new repertoire generated in this study (**S3 Text**). Finally, the expression profiles of co-expressed SSP genes were used as a template for the identification of waves of genes displaying a high level of co-regulation (R-squared > 0.80) an exemplifying specific pathogenic strategies (**Fig 3**).

**Fig 3.**
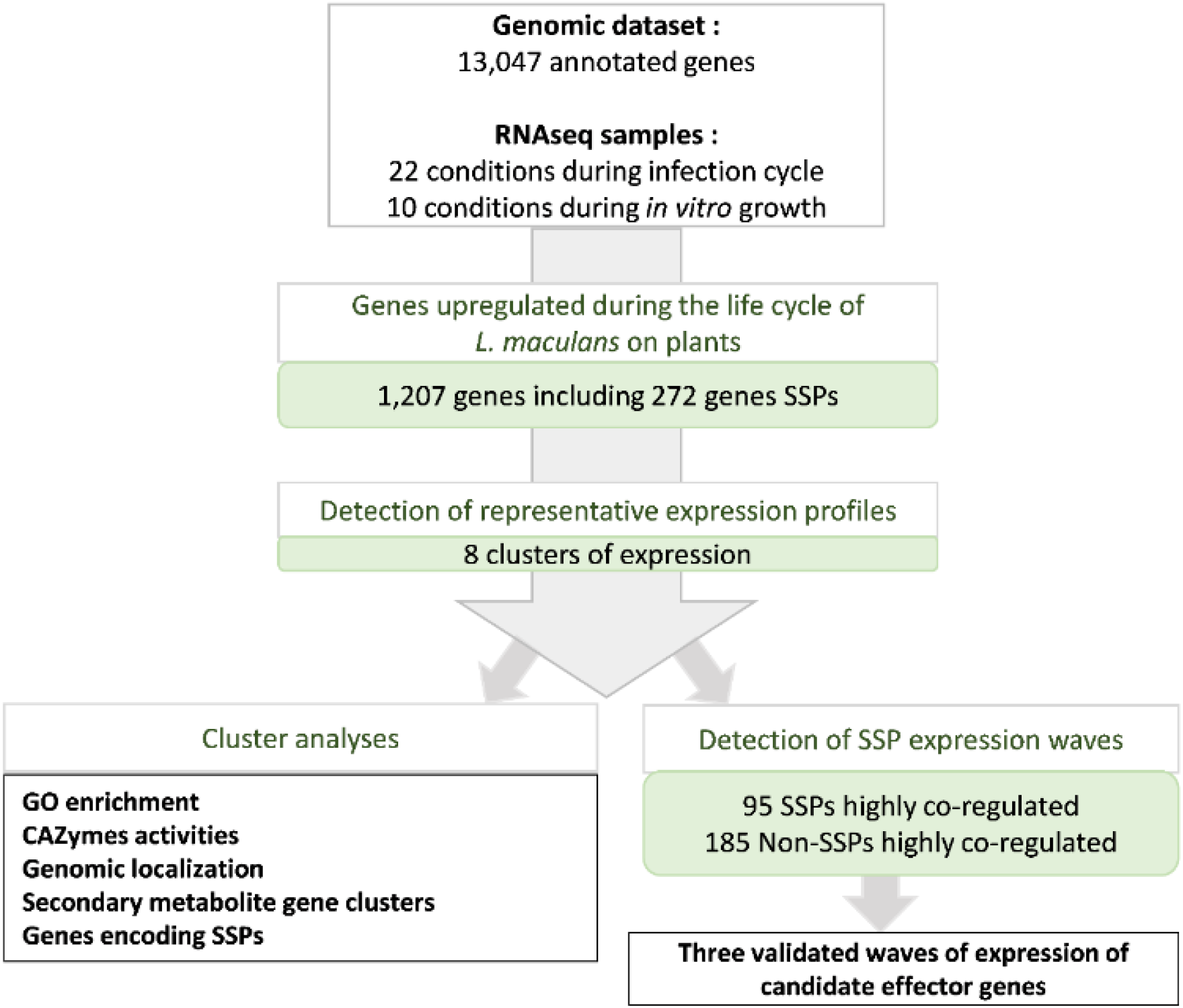
Pipeline used to describe *Leptosphaeria maculans* genes overexpressed in at least one condition *in planta*. We found that 1,207 of the 13,047 annotated genes were overexpressed in at least one of the 22 sets of conditions *in planta* relative to the 10 sets of *in vitro* conditions. These genes included 272 predicted to encode small secreted proteins (SSPs). We analyzed the set of 1,207 genes to identify the major expression profiles by gene expression clustering. We defined eight expression clusters (clusters 1 to 8). For each cluster, we analyzed expression patterns, functional enrichment, and the genomic localization of the genes. For the specific detection of highly coregulated waves of SSPs, we performed a linear regression analysis of the mean expression value for each SSP subset present in each cluster against the expression of all 13,047 genes. This analysis identified 95 SSP genes and 185 non-SSP genes as highly coregulated genes.

### Genes overexpressed during interaction with the plant define eight biologically relevant expression clusters

Relative to axenic growth conditions, 1207 genes were found to be overexpressed in at least one set of conditions *in planta*; thus, 9.2% of *L. maculans* genes are specifically mobilized during the interaction of the fungus with its host, alive or dead. Eight expression profiles were identified, referred to hereafter as clusters 1 to 8, defining a complex landscape of expression profiles (**Fig 4**). We analyzed the enrichment of each cluster in GO terms, focusing on CAZymes (Carbohydrate-active enzymes) associated with degradation activities, to identify the functions and pathways associated with each expression profile (**S2-S5 Fig**).

**Fig 4.**
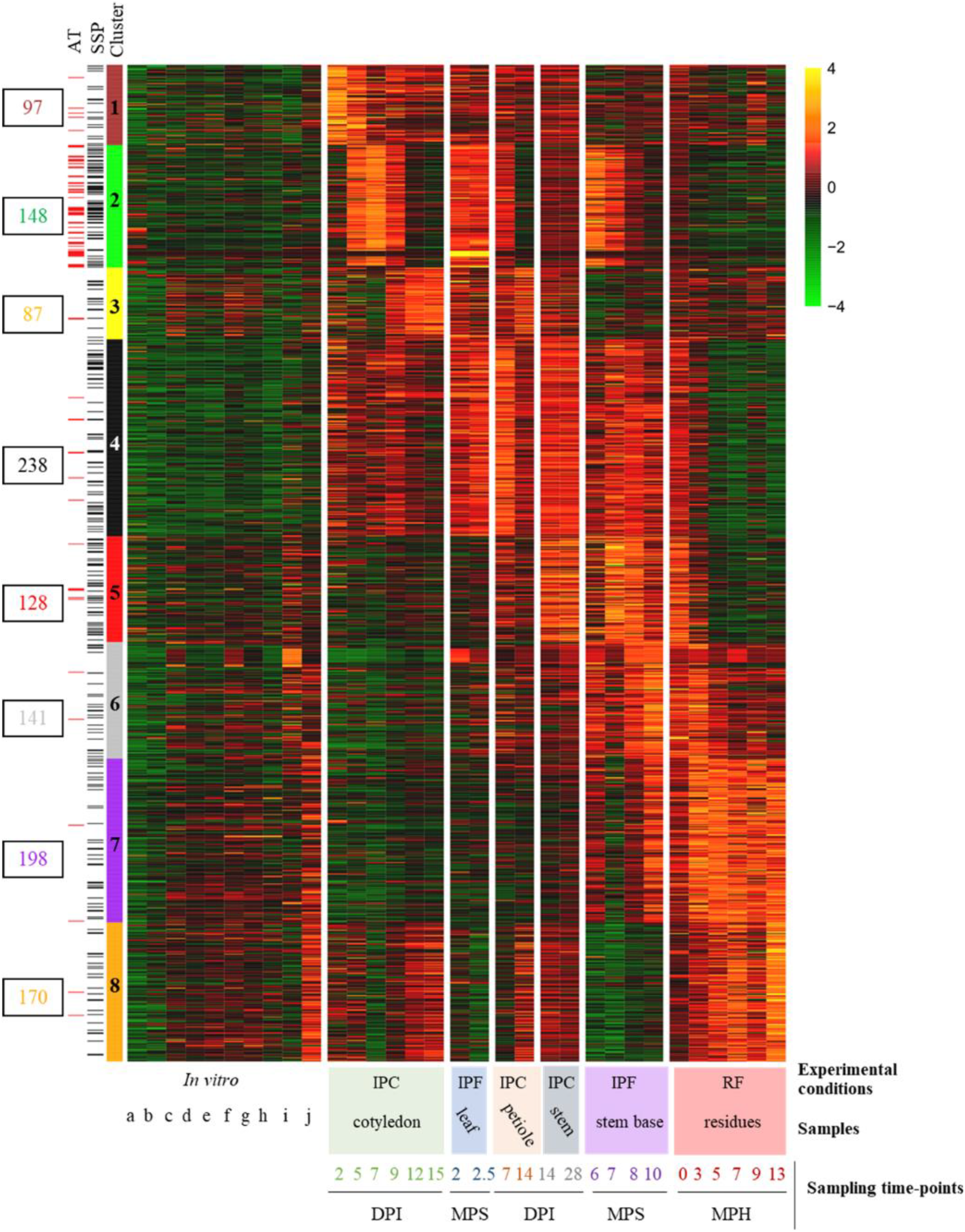
Heatmap obtained by clustering the 1,207 *Leptosphaeria maculans* genes upregulated *in planta*. For each set of conditions, we calculated the mean FPKM value of the replicates, which we then subjected to Log_2_ transformation and scaling. The self-organizing map method was used on the Log_2_(FPKM+1) values, to define eight expression clusters (clusters 1 to 8) for the 1,207 genes. The level of expression ranged from 4 (yellow) to −4 (green). Genes located in AT-rich regions (AT) or encoding small secreted proteins (SSPs) are indicated by red and black lines, respectively, on the left. The conditions are described in the legend to the *x* axis of the heatmap. Three sample features are described (i) the experimental conditions: IPF, *in planta* field conditions; IPC, *in planta* controlled conditions; RF, residues in field conditions, (ii) the type of plant tissue sampled and (iii) the sampling time points (DPI, days post inoculation; MPS, months post sowing; MPH, months post harvest. The ten sets of *in vitro* conditions are ordered as follows: conidia (a. non-germinating, b. germinating); incompatible mating conditions (c. JN2 x JN2 7days (d), d. JN2 x JN2 20d, e. JN2 x JN2 35d); compatible mating conditions (f. JN2 x Nz-T4 7d, g. JN2 x Nzt4 20d, h. JN2 x Nz-T4 35d); mycelial growth (i. in liquid medium, j. on solid medium).

Clusters 1, 2 and 3 included genes expressed sequentially during cotyledon infection and were representative of three consecutive stages occurring during the two-week period of cotyledon colonization by *L. maculans*. Cluster 1 encompassed 97 genes highly expressed at the first stages of cotyledon infection, when the conidia germinate and the hyphae penetrate the wounded plant tissues (2-5 DPI). It was enriched in GO terms corresponding to carbohydrate metabolic processes (*p* = 1×10^−2^) and catalytic activities (*p* = 1×10^−3^), including peptidase and hydrolase activities (all *p* ≤ 1×10^−2^). An enrichment in cutinase activity (*p* = 1×10^−2^), specific to this cluster, was also observed (**S2-S3, S5 Figs**).

The 148 genes of cluster 2 displayed peaks of expression during all stages of asymptomatic growth within plant tissues (cotyledon colonization 5-9 DPI, first stage of petiole colonization, first two dates of stem colonization in the field). The genes of this cluster were annotated with few GO terms or functions, but were strongly enriched in SSP genes (*p* = 2×10^−16^; **S3 Fig**). Cluster 3 encompassed 87 genes highly expressed during the necrotrophic stage on both cotyledons (9-15 DPI) and petioles, but not during the necrotrophic stem canker stage in the field. This cluster was enriched in carboxylic ester hydrolase activities (*p* = 3×10^−3^) suggesting a role in degradation activities specifically targeting the plant cell wall.

Cluster 4 encompassed 238 genes highly expressed during the shift from biotrophy to necrotrophy in cotyledons (7-9 DPI) and in stem bases infected in the field (7-8 months post-sowing, ‘MPS’). This cluster displayed the strongest enrichment in catalytic processes (*p* = 2×10^−10^) and hydrolase activities (*p* = 1×10^−12^), and a specific enrichment in proteolysis processes (*p* = 2×10^−2^; **S3 Fig**). It also displayed the largest number of CAZymes (63 genes; **S5 Fig**) targeting different substrates, mostly plant cell-wall components, such as pectin, arabinose and galactan (**S5 Fig**).

Two clusters (cluster 5 and 6) displayed an increase in expression late in stem infection, but differed in terms of the timing of expression and the functions involved. Cluster 5 contained 128 genes highly expressed during the asymptomatic colonization of stems in the field (at 7-8 MPS) and on residues immediately after harvest. It was enriched in catalytic activities (*p* = 1×10^−3^), mostly linked to saccharide degradation, suggesting involvement in activities relating to the uptake of sugar resources (carbon-oxygen lyase activity acting on polysaccharides; hydrolase activity; cellulose binding; **S3 Fig**). Cluster 6 contained 141 genes highly expressed during colonization of the stem base in the field, with maximal expression at 10 MPS, corresponding to the time at which stem canker develops. This cluster was enriched in catalytic activities (*p* = 2×10^−4^) but was also linked to cofactor binding (*p* = 3×10^−6^), monooxygenase (*p* = 1×10^−2^), and oxidoreduction activities (*p* = 5×10^−5^), involved in detoxification processes (**S3 and S4 Figs**). The higher proportion of xylanase genes in cluster 6 than in cluster 5 (**S5 Fig**) suggests a change in the mode of nutrition during the necrotrophic stage on stems.

Finally, clusters 7 and 8 were specific to saprophytic behavior on stem residues. The 198 genes in cluster 7 showed displayed an increase in expression from the last sampling on necrotic stems until the last sampling on stem residues one year after harvest. This cluster was enriched in genes relating to mono-oxygenase activities (*p* = 1×10^−4^) and contained the largest number of genes encoding LPMOs (lytic polysaccharide monooxygenases), cellulases and xylanases. Cluster 8 encompassed 170 genes specifically expressed during all stages occurring on senescent or dead plant tissue, with the highest levels of expression during saprophytic growth on residues. It was also enriched in oxidoreductase processes (*p* = 1×10^−3^), in response to oxidative stresses and oxidant detoxification associated with an enrichment in catalase (*p* = 1×10^−3^) and peroxidase (*p* = 3×10^−3^).

The different clusters of gene expression thus can be designated as follows: “Penetration and establishment” (cluster 1), “Biotrophy” (cluster 2), “Cotyledon and petiole necrotrophy” (cluster 3), “Biotrophy to necrotrophy transition” (cluster 4), “Stem biotrophy” (cluster 5), “Stem necrotrophy” (cluster 6), “Stem canker and saprophytism” (cluster 7) and “Saprophytism” (cluster 8).

### Contribution of candidate pathogenicity genes to fungal life in interaction with the plant

#### 1- Genes involved in fungal pathogenicity in other fungal species

Several genes previously shown to be involved in fungal pathogenicity were found in one of the eight expression clusters, and were mostly associated with biotrophic stages of the interaction. One RALF (rapid alkalinization factor) gene known to favor fungal infection [39] was identified in cluster 1 (Lmb_jn3_05329). Two of the six salicylate hydroxylase genes, known to compromise plant salicylic acid signaling in other models [40], were found in clusters 1 (Lmb_jn3_12582) and 2 (Lmb_jn3_11892). Three of the four LysM-domain containing genes of the genome, known to protect chitin from degradation by plant enzymes or to scavenge chitin oligomers [41], were found in clusters 2 (two genes, Lmb_jn3_00461, Lmb_jn3_10047) and 6 (one gene, Lmb_jn3_04300). Three isochorismatase genes (Lmb_jn3_11641, Lmb_jn3_10045, Lmb_jn3_06552), interfering with plant salicylate and jasmonate defense signaling in other phytopathogens, were present in clusters 1, 4 and 6, respectively. Finally, one of the two genes encoding NEP (necrosis and ethylene-inducing protein), frequently associated with necrotrophic behavior [42], was found to be overexpressed *in planta*, and belonged to cluster 4 (Lmb_jn3_10035).

#### 2- Secondary metabolite gene clusters

The genome of *L. maculans* contains 22 genes predicted by SMURF tools to encode secondary metabolite enzymes (SM) and annotated as “polyketide synthase” (PKS, 11) “non-ribosomal peptide synthetase” (NRPS, 5), or “others” (dimethylallyltryptophan synthase, 1; NRPS-like, 4; PKS-like, 1). Seven of the 1207 genes in the eight expression clusters encoded SM enzymes from the PKS or NRPS family. Four of these genes were highly expressed during necrotrophic stages of the life-cycle within the stem (cluster 6), whereas the other three were associated with clusters 2, 4 and 5 (**S3 Table**).

Among the SM genes analyzed in previous studies [27,43–44], the PKS gene involved in abscisic acid (ABA) synthesis was linked to biotrophic behavior (cluster 2), whereas genes involved in the synthesis of the toxins sirodesmin PL and phomenoic acid were associated with stem necrotrophy, together with one other PKS and oneNRPS-like gene (cluster 6) (**S3 Table**). Secondary metabolite enzymes encode an SM backbone that may be modified by tailoring enzymes, forming complex secondary metabolite gene clusters (SMGC). Here, only three of the seven SM enzyme genes overexpressed *in planta* showed co-regulation with all or part of surrounding genes of their SMGC: (i) the PKS responsible for ABA production (all genes of the ABA SMGC being included in cluster 2), (ii) the NRPS responsible for sirodesmin PL production (16 of the 25 genes of the SMGC included in cluster 6), and (iii) an unknown NRPS-like gene, co-expressed with nine other surrounding genes also included in cluster 6 (**S6 Fig**).

#### 3- Genes involved in *L. maculans* pathogenicity

Thirty-six of the 52 genes previously identified as candidate pathogenicity genes in *L. maculans* (**S4 Table**) did not belong to the SM or avirulence effector gene (see below) categories. Fifteen of these genes were identified within a single cluster (**S4 Table**): cluster 1 (2 genes), cluster 2 (5 genes), cluster 3 (1 gene), cluster 4 (4 genes), cluster 5 (2 genes) or cluster 7 (1 gene). Nine had previously been selected as putative pathogenicity genes on the basis of their overexpression during infection, for inactivation with CRISP-cas9, but the inactivation of these genes individually resulted in no visible pathogenicity defect at the cotyledon stage [45] (**S4 Table**). Only one gene, encoding a 3-ketoacyl-thiolase (cluster 1), was shown to be involved in pathogenicity, but the mutant also displayed impaired *in vitro* growth and germination [28] (**S4 Table**). The other five genes have not been validated by functional approaches and are, thus, still considered to be candidate pathogenicity-related genes. Four of these genes (three GH genes and one ABC transporter) were included in the late expression clusters 4, 5 and 7, suggesting that they may be *bona fide* pathogenicity genes (**S4 Table**).

By contrast, another 21 putative pathogenicity genes did not belong to any of the expression clusters. Nineteen had previously been analyzed in functional studies (knockout or silenced mutants), and 10 were shown to induce a loss or reduction of pathogenicity following inactivation (**S4 Table**). However, the inactivation of six of these genes also induced defects in growth, sporulation or germination [30,33,36,45–46], suggesting a general effect on different stages of the fungal life-cycle with no specific overexpression during interactions with the plant.

#### 4- Validated or candidate SSP effectors of *L. maculans*

SSPs have been identified in *L. maculans* and were classified as “early” effectors (including all known avirulence proteins, inducing an avirulence phenotype when matching the corresponding resistance genes) or “late” candidate effectors by Gervais *et al*. [14]. The role of avirulence proteins as effectors suppressing plant defense responses or increasing/decreasing the size of cotyledon lesions has been investigated in a few cases following comparisons of near-isogenic *L. maculans* isolates, gene silencing or complementation (**S4 Table**, [21–22,47–48]). The most recently published version of the genome assembly of *L. maculans* isolate JN3 ([49]; **S4 Text**) included a number of major changes. We therefore produced a new repertoire of genes encoding putative SSPs, encompassing 1,070 genes (8.2% of the entire *L. maculans* gene set) (**S3 Text**, **S7 Fig, S5 and S6 Tables**).

All expression clusters displayed a significant enrichment in genes encoding SSPs (**S3 Fig**). We found that 272 of the 1207 genes upregulated during the *L. maculans* interaction with *B. napus* (22.5%) encoded SSPs. This enrichment was limited in clusters 1, 3, 6, 7 and 8, which contained only 14-17% of SSP-encoding genes (0.013 < *p* < 3.10^-5^), but three other clusters displayed a much higher level of enrichment in genes encoding SSPs: cluster 4 (23.1%, *p* = 7×10^−16^), cluster 5 (31.2%, *p* = 2×10^−16^) and cluster 2 (45.3%, *p* = 2×10^−16^). Interestingly, both clusters 2 and 5 are associated with biotrophic behavior of the fungus. All the currently known *AvrLm* genes belonged to the “Biotrophy” cluster (cluster 2) (**S4 Table**). This cluster displayed sequential rounds of overexpression/repression, with three peaks of overexpression occurring during the three biotrophic stages (5-9 DPI on cotyledons, leaf infection in the field; 7 DPI on petioles; 6-7 MPS in field plant stems) (**Fig 5; S8 Fig**).

**Fig 5.**
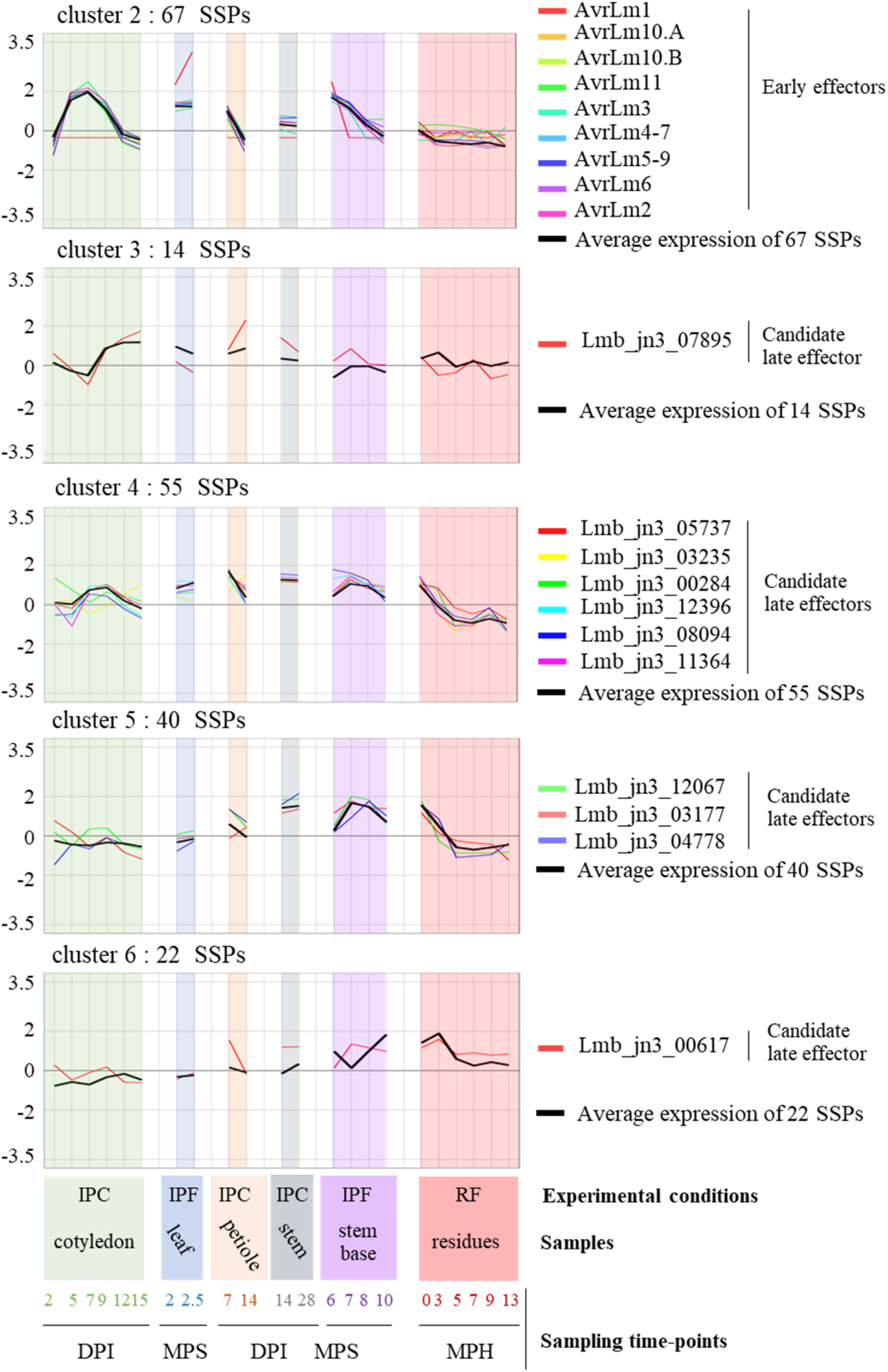
Expression of known and putative effectors of *Leptosphaeria maculans* within the eight expression clusters. The scaled Log_2_(FPKM +1) expression values for the nine avirulence effector (*AvrLm)* genes, and the eleven candidate late effector genes shown to be upregulated in at least one set of conditions *in planta* are represented and grouped according to cluster assignment. The mean scaled Log_2_(FPKM +1) values of all the small secreted protein (SSP) genes in each cluster are indicated (black bold curve). The total number of SSP genes in each cluster is indicated. The conditions are described in the legend to the *x* axis. Three sample features are described: (i) the experimental conditions: IPF, *in planta* field conditions; IPC, *in planta* controlled conditions; RF, residues in field conditions. (ii) the type of plant tissue sampled and (iii) the sampling time points (DPI, days post inoculation; MPS, months post sowing; MPH, months post harvest).

The ten “late” effectors, expressed during stem colonization in controlled conditions were scattered over four different clusters (3, 4, 5, 6), along with a series of other genes encoding SSPs (cluster 3, 14 SSPs; cluster 4, 55 SSPs; cluster 5, 40 SSPs and cluster 6, 22 SSPs) (**Fig 5**). The “late” effectors in clusters 5 and 6 were specific to stem colonization (and the first stages of saprotrophy on residues), but those in clusters 3 and 4 were also expressed during cotyledon infection. These “late” effector candidates were associated with either biotrophic behavior on stems (cluster 5) or necrotrophic behavior on cotyledons and stems (clusters 3 and 6).

Clusters 1, 7 and 8, were also found to be enriched in genes encoding SSPs (*p* value < 5×10^−3^). They contained 16, 33 and 24 SSP genes, respectively. Candidate effectors may, therefore, be mobilized very early in colonization (cluster 1) and during saprophytic life (clusters 7 and 8) (**S8 Fig**).

Additional RNA-Seq samples (ascospores germinating on unwounded cotyledons 24 and 48 hours post inoculation; stem base samples 2, 3 and 5 MPS) were available. They provided too few *L. maculans* reads for statistical analyses, but these samples allowed us to confirm that genes belonging to cluster 2, and “early” effector genes in particular, were among the genes most strongly expressed both during ascospore germination on cotyledons and very early stages of stem colonization in winter (**S5 Text**, **S7 Table**).

### Histone methylation associated with gene regulation

All cloned avirulence genes of *L. maculans* identified to date are associated with H3K9me3. Furthermore, H3K9me3 and H3K27me3 domains are significantly enriched in genes encoding effectors, regardless of their expression [38]. During axenic culture, 7,373 *L. maculans* genes are associated with H3K4me2, 104 with H3K9me3, 2,020 with H3K27me3 and 101 with the two heterochromatin histone modifications [38]. We combined the genome-wide histone maps previously generated *in vitro* with our analysis of gene expression during infection, to determine whether (i) all genes, (ii) only (putative) effector genes, or (iii) only certain subsets of genes upregulated *in planta* were associated with a particular chromatin landscape. H3K4me2-domains encompass 56% of *L. maculans* genes, but only 20% of the genes upregulated *in planta* were associated with H3K4me2 (◻^2^ test=662; *p* < 2×10^−16^). By contrast, the genes upregulated *in planta* were significantly enriched in genes located in an H3K27me3 domain, as 44% of the upregulated genes were located in such a domain (◻^2^ test = 749, *p* < 2×10^−16^). Strikingly, only 205 genes were located in H3K9me3 or H3K27me3 domains *in vitro*, but 85 of these genes were upregulated *in planta*. Thus, regardless of the wave of expression concerned, the genes upregulated *in planta* were enriched in genes associated with heterochromatin *in vitro*. Only cluster 2 was significantly enriched in genes associated with the H3K9me3 histone modification (**Fig 6**). Interestingly, all clusters of genes upregulated *in planta* relative to axenic culture were enriched in H3K27me3-associated genes *in vitro,* whereas these clusters contained significantly fewer genes located in H3K4me2-domains *in vitro* than the rest of the genome (**Fig 6**). The integration of RNA-seq and ChIP-seq data revealed that the genes upregulated *in planta* were not randomly located in the genome of *L. maculans*, with H3K4me2-domains significantly depleted of genes upregulated during rapeseed infection, and both types of heterochromatin domains significantly enriched in such genes.

**Fig 6.**
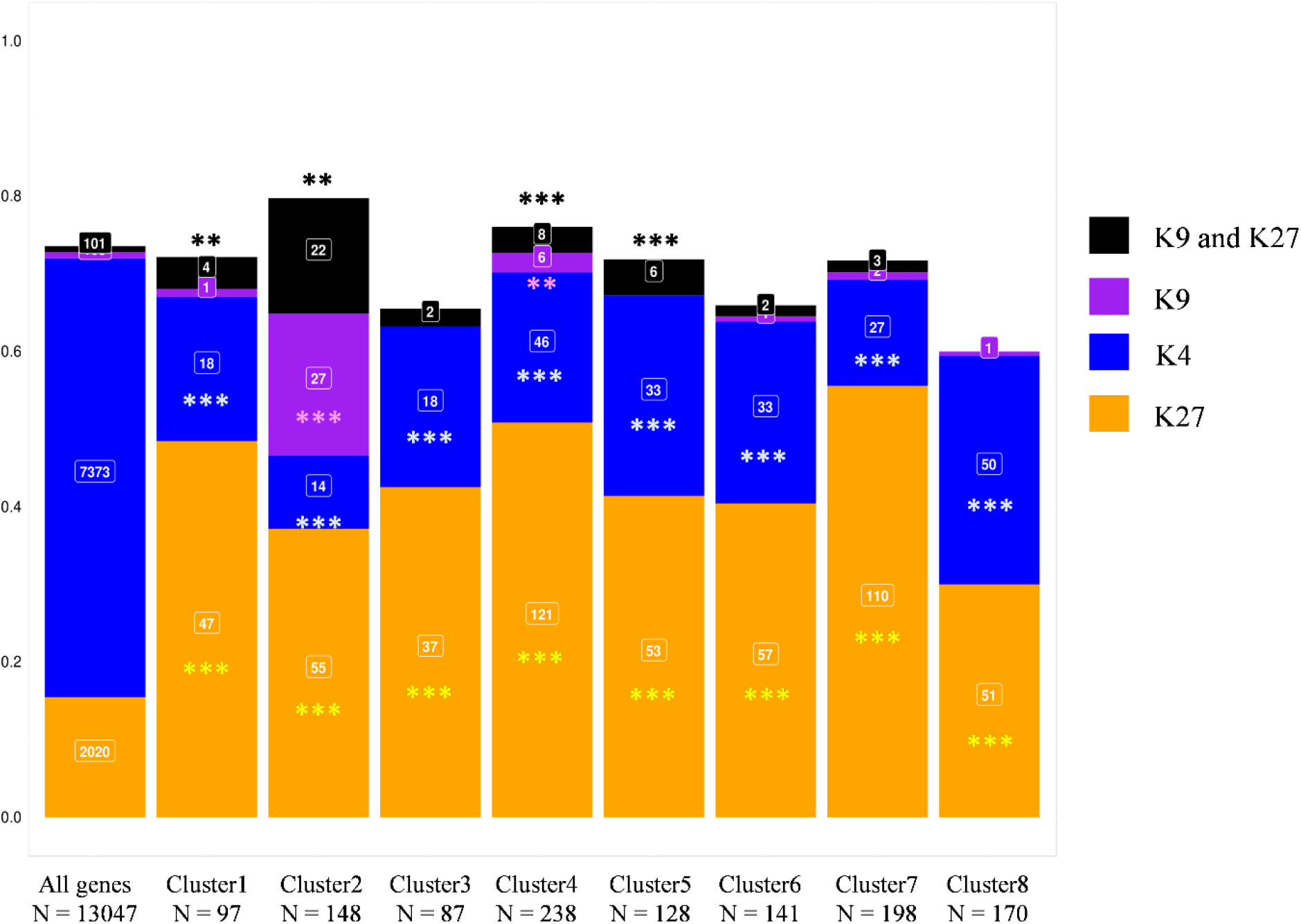
Association of *Leptosphaeria maculans* genes upregulated *in planta* with the histone modification marks. The genome-wide histone map was generated by ChIP-seq analysis during axenic growth [38]. The proportion of genes associated with any of the histone modifications H3K4me2, H3K9me3 and H3K27me3 throughout the entire genome was used as a reference set for detecting the over- or under-representation of specific methylation marks in the eight gene clusters overexpressed *in planta*, in chi-squared tests (***: *p* < 0.001, **: *p* < 0.01). K4, H3K4me2; K9, H3K9me3; K27, H3K27me3.

### The co-expression of effector genes defines waves of co-regulated gene expression

Almost all the expression clusters were enriched in genes encoding SSPs. However within each cluster, the genes encoding SSPs could have different expression profiles, due to the clustering method (**S8 Fig**). We aimed here to increase the robustness of the detection of SSP expression waves. We defined eight SSP reference expression profiles by calculating the mean levels of expression for SSP genes within the eight clusters. We selected genes displaying significant co-regulation (SSP and non-SSP genes), by analyzing the linear regression between the expression patterns of the entire set of 13,047 genes and that of each SSP reference expression profile. Genes significantly co-expressed with the reference expression wave tested (R^2^>0.80, *p* <0.05) were included in the new SSP expression waves and were considered to be “highly coregulated genes”.

For clusters 1, 3, and 8, no highly coregulated genes could be identified (**Fig. 7**). For the other five clusters, waves of highly coregulated genes were identified, each containing five to 39 highly coregulated SSP genes (including 90 SSP genes from the 178 previously assigned to these clusters), together with 186 highly coregulated genes encoding other types of proteins (**Fig 7**). Two waves of expression, corresponding to clusters 5 and 6, displayed coregulation of a limited number of SSP genes (seven and five, respectively, i.e. 17% of the SSP genes of these clusters), together with 20 and 25 genes encoding other proteins, respectively. However, only 15 (cluster 5) and 13 (cluster 6) of these genes belonged to the clusters (**Fig 7**), suggesting that additional coregulated genes are either expressed in axenic conditions, or do not display specific regulation during plant infection. For clusters 2, 4 and 7, a higher proportion of highly coregulated SSP genes was found, with 58% (exact confidence interval [45-70%]), 34% ([22-48%]) and 63% ([45-79%]) of the SSP genes of the clusters retained in the waves of highly coregulated genes, respectively (**Fig 7**). Thus, clusters 2 and 7 displayed the highest proportion of highly coregulated SSP genes, suggesting that the regulation of expression is much tighter in these two clusters than in the others. The highly coregulated genes in the “Biotrophy” wave (contained in cluster 2) included 39 SSPs, 23 (59%) of which displayed no sequence matches to the NCBI nr protein database (**S8 Table**) and could be considered *L. maculans*–specific effectors. These genes also included other genes mentioned above, such as the genes of the ABA cluster, providing addition support for their role in the biotrophic parts of the interaction. The highly coregulated genes in the “Stem canker and necrotrophy” wave included most of the SSP genes of the cluster, but only two of these genes (9.5%) were specific to *L. maculans*. This wave also included a large number of genes (24) encoding CAZymes.

**Fig 7.**
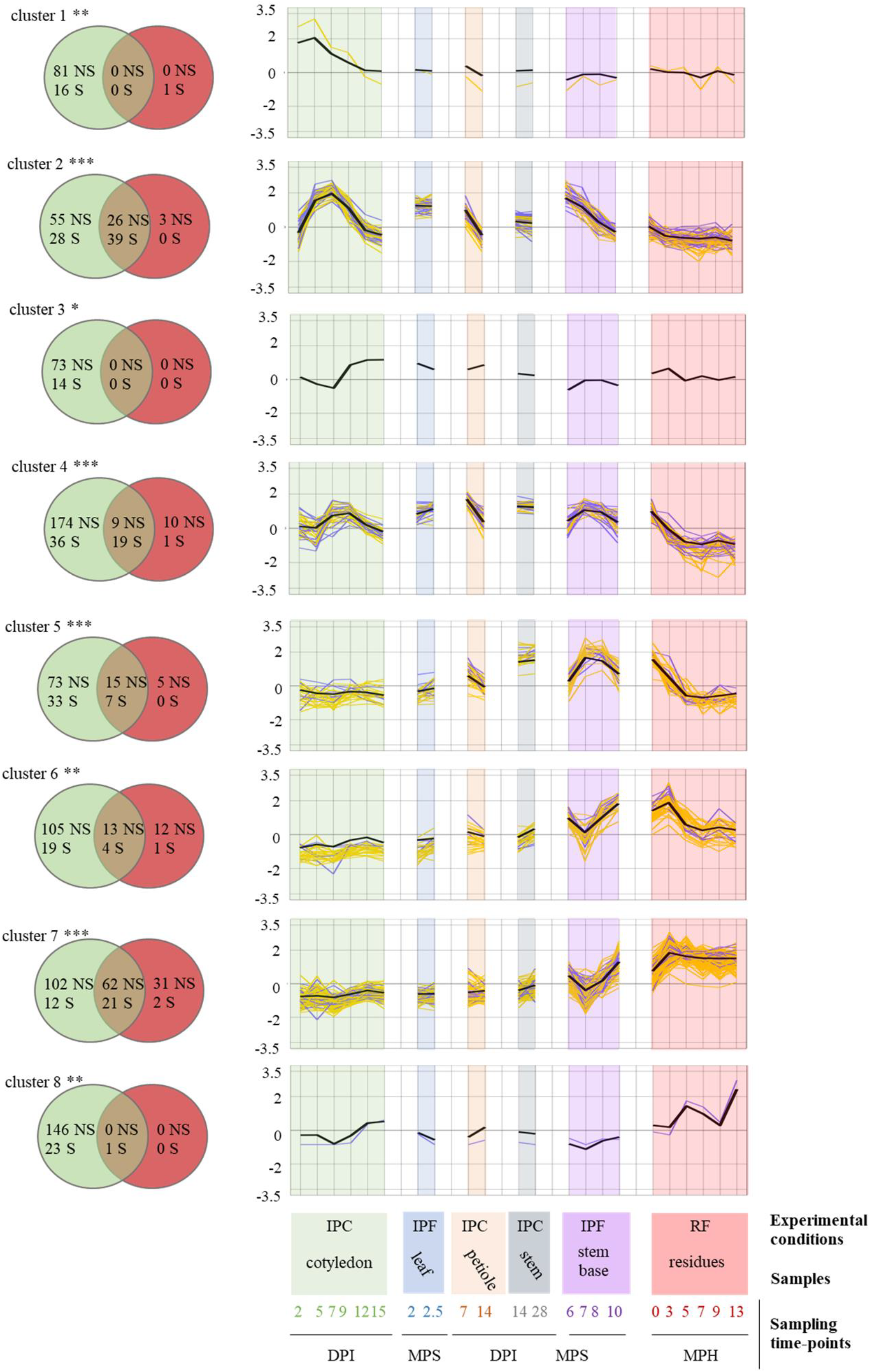
Identification of highly coregulated *Leptosphaeria maculans* genes based on mean SSP gene expression *in planta*. The mean scaled Log_2_(FPKM +1) values for small secreted protein (SSP) genes in each cluster (cluster 1 to 8) was used to create the eight reference expression profiles (black bold curve). The 22 sets of biological conditions used to average SSP gene expression are described in the legend to the *x* axis. Three sample features are described: (i) the experimental conditions: IPF, *in planta* field conditions; IPC, *in planta* controlled conditions; RF, residues in field conditions (ii) the type of plant tissue sampled and (iii) the sampling time points (DPI, days post inoculation; MPS, months post sowing; MPH, months post harvest. A linear regression analysis was performed to determine the correlation between the scaled Log_2_(FPKM+1) value for the whole set of genes in the 22 sets of conditions and these eight reference expression profiles. A gene was considered to be highly correlated if its expression fitted the reference expression profile with an R-squared value > 0.80 and a corrected *p*-value < 0.05. The expression values of the resulting highly correlated gene subsets (SSP in blue and non-SSP in yellow) are plotted. The Venn diagrams show the number of genes (S: SSP, NS: Non-SSP) present in each cluster (green circles) and the number of genes identified as highly correlated with the corresponding expression profile (red circles). The asterisks indicate the significance of the SSP enrichment in the initial cluster (***: *p* < 0.001, **: *p* < 0.01).

## Discussion

In this paper, we used large-scale transcriptomic analyses to decipher the complex life cycle of a phytopathogenic fungus, to identify genes paramount for its interactions with the host plant and to shed light on the underlying regulatory mechanisms. In particular, given that many parts of the *L. maculans* life cycle remain poorly described, we focused on how the fungus copes with its different lifestyles, their timing and the transitions between them, and the regulation of gene expression throughout the long pathogenic/saprophytic life of the fungus. We generated RNA-Seq data for biological samples corresponding to the main relevant stages of infection in controlled conditions or in field conditions, and at different time scales (from days to years). Using this large dataset, we then focused on sets of genes mobilized exclusively when the fungus interacts with the plant. We used 10 different sets of axenic growth conditions promoting different aspects of the fungal life-cycle (e.g. vegetative growth, asexual sporulation, sexual reproduction) as basic expression features for the identification of pathogenicity genes *per se*, contrasting with the common practice of using only a few axenic conditions as controls [7,13–14] or performing pairwise comparisons of the infection time course [11]. We dealt with the variable number of fungal reads between samples and the diverse *in vitro* conditions, by applying very stringent conditions to determine whether a gene was specifically overexpressed (LogFC>4, *p* < 0.01). With these criteria, we showed that 1,207 genes (less than 10% of the genes of the fungal genome) were overexpressed during interactions with the living or dead plant relative to axenic conditions. These genes may be considered the maximal set of genes mobilized by the fungus to infect the plant without involvement in basal biological processes. These 1,207 upregulated genes were consistently grouped into eight expression profiles, some of which displayed strong coregulation of a number of the genes within the cluster. The sequential expression profiles are consistent with the associated pathogenicity functions, confirming the relevance of this RNA-Seq study performed over a large timescale.

### Transcriptome-based description of a complex fungal lifecycle

Our RNA-Seq data highlighted and strengthened our knowledge of the biology of the fungus all along its life-cycle, including for “obscure parts” of this cycle, as follows (**Fig. 8**) :

i. Ascospores, which are difficult to generate for pathogenicity tests in controlled conditions, are known to be the main source of infection in the field, germinating within a few hours to produce a hypha that penetrates leaf tissues via stomata or wounds [50]. We found that germinating hyphae very rapidly (2 DPI) produced “early” effectors (including known AvrLm effectors), detected only 5-7 DPI as a massive peak of expression when inoculations were performed with conidia ([11,13–14], this study). This result is consistent with microscopy-assisted RNA-Seq analyses in other models, which have suggested that a few hyphal tips produce early effectors to counter the initial defense reactions of the plant upon contact [4,51]. This suggests that, like other plant-associated fungi, *L. maculans* manipulates the plant as soon as it enters the tissues, to initiate its own growth within plant tissues, without causing symptoms.
ii. Following the infection of leaves or cotyledons and primary leaf spots development, the hyphae grow within the intercellular spaces within tissues in advance of the leaf spots towards and within the petiole and then enter the stem tissues [52–54]. Our RNA-Seq data highlight the kinetics of this process and the lifestyle transitions occurring during cotyledon infection, with three successive waves of gene expression related to different stages of infection (penetration, biotrophy, and necrotrophy). Consistent with the biological data, the pattern of gene expression during petiole colonization was similar to that in cotyledons, with a symptomless stage during which the same set of genes was mobilized in cotyledons or petioles (**Fig 8**).
iii. We also found that isolates infecting leaves in the field had a pattern of gene expression similar to that of a reference isolate used to inoculate cotyledons in controlled conditions at 7 and 9 DPI, demonstrating that infections reproduced with routine protocols in controlled conditions can accurately mimic some stages of field infection.
iv. The monitoring of stem samples for a full year in field conditions provided new information about the dynamics of stem colonization and necrosis in the field. We found that the fungus was present very early in the stem base tissues, with fungal reads detectable as early as December, as little as one to two months after the initial leaf infection. However, levels of fungal transcriptomic activity remained very low between December and February, and it was only at the end of winter (from March on) that the fungus displayed a quantifiable, steady increase in transcriptomic activity.
v. Fungal life within stem tissues, over a period of eight months, was characterized by complex patterns of gene expression, with some of the genes concerned expressed exclusively during stem colonization, highlighting the lengthy transition from biotrophy to necrotrophy leading to stem canker (**Fig 8**). The gene expression patterns observed for stem infection at six and seven months post sowing (March-April) were very similar to those observed for biotrophic stem infection in controlled conditions at 14 and 28 DPI, again demonstrating that controlled conditions adequately reproduce at least one stage of stem colonization in the field.
vi. The fungus has been reported to survive on residues for between two and four years, depending on the climatic conditions [50]. However, we currently know nothing about how *L. maculans* survives as a saprobe on residues. A recent metabarcoding study showed that *L. maculans* was the predominant fungal species on rapeseed residues, on which many other fungal species, including other phytopathogens were also present, highlighting complex interplay between the species [55]. Our RNA-Seq analysis of residues left on the soil for one year confirmed these findings and systematically revealed transcriptomic activity of *L. maculans* on the residues, with a mean of 22% of the total RNA-Seq reads corresponding to *L. maculans* throughout the year, but with seasonal variations.

**Fig 8:**
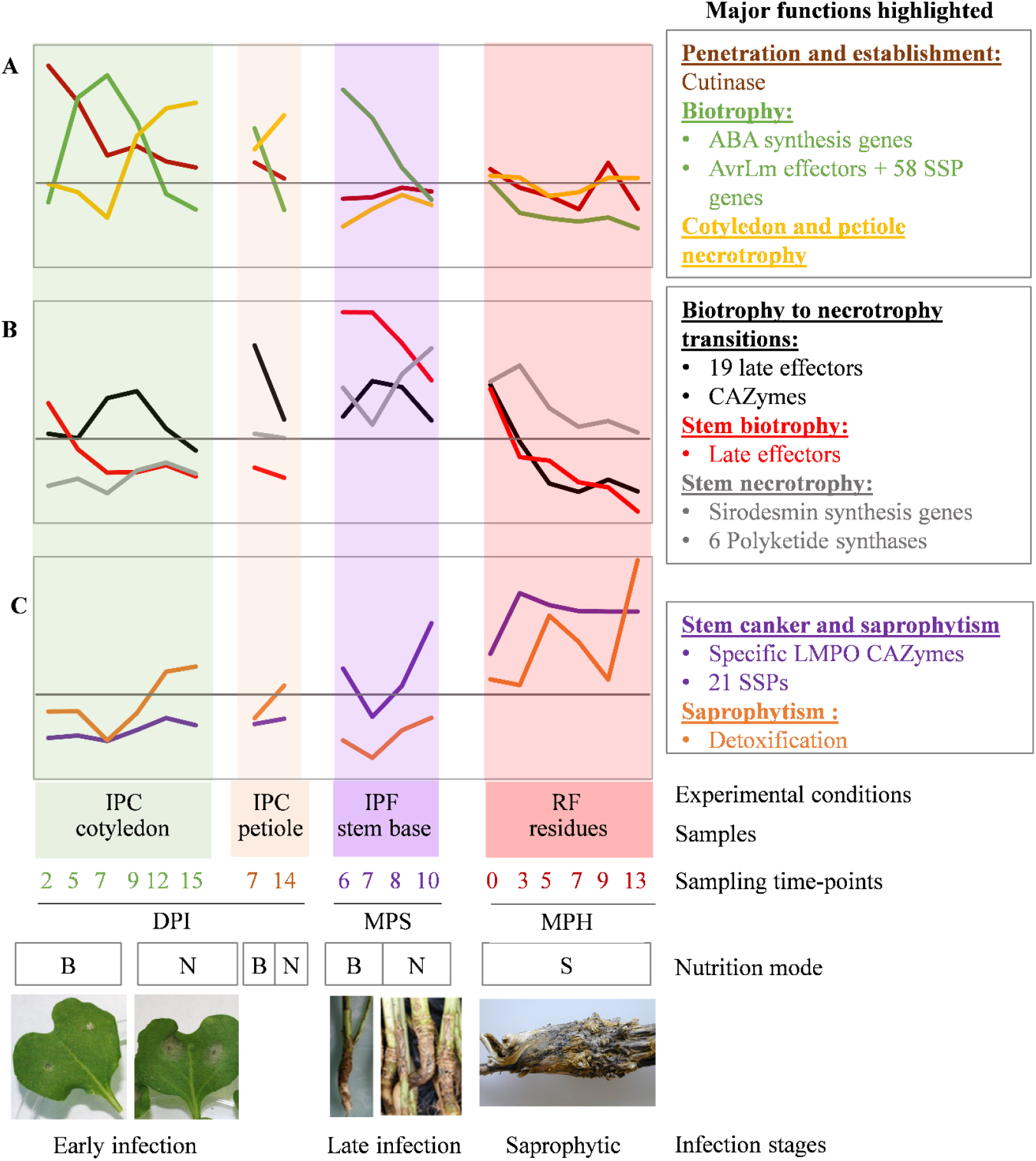
Overview of the eight expression profiles detected during the infection cycle of *Leptosphaeria maculans*. Each curve represents the mean expression value for the genes making up the eight clusters described in this study. The eight profiles are divided into three different plots, depending on the timing of their major expression peaks: (A) early infection and colonization of cotyledons, (B) late colonization of stems, (C) development on crop residues. The characteristics associated with each sample are indicated: mode of nutrition (B: biotrophy, N: necrotrophy, S: saprotrophy); the experimental conditions: (i) IPF, *in planta* field conditions; IPC, *in planta* controlled conditions; RF, residues in field conditions. (ii) the type of plant tissue sampled and (iii) the sampling time points (DPI, days post inoculation; MPS, months post sowing; MPH, months post harvest. We did not include field-infected leaves and the stem infected in controlled conditions, to prevent redundancy and generate a simplified model. The major gene functions identified in each gene cluster are shown on the right.

### Expression profiles and characteristics of genes specifically involved in interaction with the plant

We observed successive or partly overlapping expression profiles, highlighting various life traits or feeding strategies specific to a particular lifestyle or colonized tissue (cluster 1, cluster 2, cluster 4, cluster 8), or with a combination of lifestyle/tissue specificities (cluster 3, cluster 5, cluster 6, cluster 7) (**Fig 8**). For example, necrotrophic behavior is tissue-specific, with genes expressed during cotyledon or petiole necrosis (cluster 3) or in stem tissues at the end of the growing season (cluster 6; **Fig 8**). By contrast, cluster 2 is involved in biotrophic behavior, whatever the tissue colonized.

The set of 1,207 genes specifically expressed during interaction with the plant is strongly enriched in candidate effectors, with enrichment in effectors observed for all expression clusters. As shown for cotyledon infection in other studies [13], and expanded here to other biotrophic or necrotrophic stages of plant infection, biotrophic behavior was associated with enrichment in very few, if any GO terms, whereas biotrophy-necrotrophy transitions were strongly enriched in genes involved in catalytic processes and CAZymes. Strict necrotrophic behavior within stems is associated with an overrepresentation of PKS-encoding genes (including the sirodesmin biosynthesis gene cluster) relative to other necrotrophic clusters.

The fungus expresses a set of 148 genes in all biotrophic/endophytic/asymptomatic stages of colonization (cluster 2). Interestingly, we found that the expression of these genes was up- and down-regulated on multiple occasions during the interaction with the plant. The recycling of the same set of genes for a similar lifestyle highlights the key role played by this set of genes in establishing symptomless growth within the plant, with at least one set of stem-specific genes (cluster 5) also associated with biotrophic behavior, but only during stem colonization. Cluster 2 (and, to a lesser extent, cluster 5) was found to be strongly enriched in genes encoding candidate effectors, which are generally thought to act by compromising plant immunity [15]. One of these candidate effectors, AvrLm10A, has been shown to compromise leaf symptom development [23], and two others, *AvrLm4-7* and *AvrLm1,* interfere with SA and ET signaling [47–48]. In addition, cluster 2 includes genes reported in various models to be involved in the suppression of plant defense responses (salicylate hydroxylase or the ABA cluster; [27,56–57]) or the scavenging of chitin oligomers, such as the LysM genes [41]. This strongly suggests that cluster 2 plays an important role in hiding the fungus from the plant surveillance machinery, or in compromising or suppressing plant defense responses to various extents.

Our findings suggest that the previous classification of effectors as “early” or “late” [14] is not relevant. “Early” effector genes were found to be expressed on several occasions during symptomless cotyledon infection and colonization, the symptomless colonization of petioles and during at least three months of symptomless growth in the tissues of the stem base. Similarly, the “late” effector genes identified by Gervais *et al*. [14] and thought to be specific to stem colonization according to experiments in controlled condition, were actually found in four different expression clusters, only one of which was specific to biotrophic behavior in stem tissues (cluster 5, **Fig 8)**.

Importantly, cluster 2 contains all the genes encoding avirulence proteins identified to date in *L. maculans*. During leaf or cotyledon infection, these avirulence proteins can be recognized by the products of the major resistance genes (*Rlm* genes) of the plant, resulting in complete resistance to avirulent isolates. The re-use of these same genes at other stages of plant colonization strongly suggests that recognition can also take place during the systemic growth of the fungus within petioles or stems, resulting in efficient resistance at these stages of colonization, provided that the *Rlm* genes are constitutively expressed in all plant tissues. In addition to the nine already cloned avirulence genes, cluster 2 contains 58 additional SSP genes (39 of which are highly coregulated with all *AvrLm* genes), which are candidates of choice for the screening of genetic resources for novel resistance genes.

We also found that effector gene expression was not restricted to interaction with the living plant. Genomic studies on fungal saprotrophs focus on CAZymes and class-II peroxidases [58]. We show here that *L. maculans* recruits a typical cocktail of CAZymes, including the lytic polysaccharide mono-oxygenases (LPMO) involved in wood decay, but also peroxidases, cytochrome P450 and stress response genes, during one year of life on crop residues. In addition, cluster 7 encompasses 21 SSP genes specifically expressed during saprophytic growth on stem residues (**Fig 8**). It has been suggested that effector genes arise from genes used by saprobes to suppress ecological competitors [59]. The steady expression of a series of coregulated candidate effector genes during many months of life on residues, regardless of the changes in the mycoflora present on the residues over time [55], may indicate a need for the fungus to produce a minimal set of molecules involved in ecological competition.

### Heterochromatin-based regulation of expression as a key player in the lifetime coregulation of genes involved in pathogenicity

One of the key findings of our work is that all clusters of genes overexpressed during interaction with the host are significantly associated with localization within a heterochromatin domain during axenic growth. Evidence is accumulating, for many other plant-associated fungi, regardless of their lifestyles and modes of nutrition, that the genes involved in fungus-plant interaction are often present in regions of heterochromatin within the genome (e.g. in [60–62]). Genes highly expressed during infection are often associated with heterochromatin during axenic culture (e.g. in *Z. tritici* or *L. maculans* [4,37,63]) and histone modifications (H3K9me3 and / or H3K27me3) have been shown to play a major role in regulating secondary metabolite gene clusters or effector genes (e.g. in *F. graminearum*, *F. fujikuroi*, *Z. tritici*, *Epichloe festucae* [64–68]). In the endophyte *E. festucae* and the pathogen *Z. tritici*, the derepression of putative effector genes or secondary metabolite gene clusters upon infection is associated with an underlying dynamic of histone modifications, which has been experimentally validated [64,67]. These data suggest that there is a generic chromatin-based control of plant infection mechanisms. We previously showed that, in *L. maculans,* heterochromatin domains (either H3K9me3 or H3K27me3) display significant enrichment in putative effector genes, and that all known avirulence genes are associated with H3K9me3 during axenic growth [38]. We found here that the genes included in the eight clusters specifically overexpressed during interaction with the plant relative to axenic growth were also enriched in H3K9me3 and/or H3K27me3 histone modifications *in vitro*. One striking finding of our study is that genes from the various waves of expression are scattered throughout the genome, but are significantly associated with heterochromatin, facilitating the temporal regulation of sets of genes involved in the same process. All the eight expression clusters identified in the lifecycle of *L. maculans* contained genes displaying significant enrichment in H3K27me3 heterochromatin *in vitro*. This histone modification is an important regulator of development and response to several biotic and abiotic stresses, in various fungal or plant models (e.g. [69–70]). By contrast, genes associated with biotrophic or biotrophic/asymptomatic life within the plants (cluster 2, **Fig 8**) displayed enrichment in H3K9me3 *in vitro*. The genes in this cluster are located in TE-rich regions of the genome and were found to be more enriched in heterochromatin modifications than the genes of the other clusters. They included the largest number of candidate effector genes, and the largest number of highly coregulated genes (65 genes). In *L. maculans*, the TE-rich regions are comprised of heterochromatin, leading to the silencing of associated genes; loss of the repressive histone modification H3K9me3 is a prerequisite to induce the expression of these genes [37]. The unique association between genes involved in the symptomless spread of the fungus and H3K9me3 suggests that this histone modification is involved in a regulatory mechanism important for the concealment of the fungus from the plants defenses. Conversely, this cluster also contains all the known avirulence genes, providing additional support for the role of H3K9me3 as the primary regulator of expression for genes at the forefront of the battle with the plant, and as a target of choice for the establishment of effector-triggered immunity during co-evolution of the plant and the pathogen.

The genes associated with H3K9me3 in cluster 2 appear to be more strictly coregulated than those in other clusters, in which genes are associated with H3K27me3. Our findings suggest that chromatin remodeling during interaction with the plant is highly dynamic, rather than occurring only once when the fungus infects the plant, as might have been anticipated from analyses of species with a short lifecycle, or from the few days of interaction amenable to laboratory experiments. This combined analysis of transcriptomic and epigenomic data thus supports the notion that the finely tuned temporal progress of fungal infections is underpinned by a highly dynamic phenomenon based on the successive opening and closing of different genomic regions.

## Materials and methods

### Plant and fungal materials

Isolate JN2 (v23.1.2) was used in all experiments. Unless otherwise stated, the fungus was cultured on V8-agar medium, and conidia for inoculation tests were produced as previously described [71]. Isolate Nz-T4, which is sexually compatible with JN2, was also used in *in vitro* crossing experiments. Both isolates were described by Balesdent *et al*. [72].

Three winter *B. napus* cultivars were used, depending on the experiment: ‘Darmor-*bzh’*, the reference sequenced cultivar [73], ‘Darmor’ and ‘Bristol’. ‘Darmor-*bzh*’ is a dwarf isogenic line resulting from the introduction of the dwarf *bzh* gene into ‘Darmor’ [74]. ‘Darmor-*bzh’* and ‘Darmor’ are characterized by a high level of quantitative resistance in the field, and ‘Bristol’ is moderately susceptible [75].

### Production of biological material for RNA sequencing

#### *In vitro* culture

The fungus was grown under different *in vitro* conditions, to mimic as many physiological stages as possible. All replicates used for RNA sequencing were true independent biological replicates. The following physiological conditions were reproduced: mycelial growth on V8-agar medium or in liquid Fries medium, pycnidiospore production on V8-agar medium, pycnidiospore germination in liquid Fries medium, and sexual mating *in vitro* at three time points (7, 20 and 35 days). All samples were immediately frozen in liquid nitrogen after recovery and stored at −80°C until RNA extraction. Detailed protocols for the production of samples in each set of conditions are provided in the S6 Text.

#### Cotyledon inoculation and sampling

The cotyledons of 10-day-old plants of cv Darmor-*bzh* were inoculated with pycnidiospore suspensions, as previously described ([76]; S6 Text). Control plants were mock-inoculated with sterile deionized water. Samples for RNA-Seq were recovered at 0 (with or without wound), 2, 5, 7, 9, 12, 15 and 17 DPI. At each time point, eight cotyledons from eight different plants were randomly selected. The plant tissues around the inoculation site were cut with a 10 mm-diameter disposable hole-punch and the 16 corresponding samples were pooled together in a Falcon tube, immediately frozen in liquid nitrogen and stored at −80°C until extraction. Two replicates were recovered at each time point and the whole experiment was repeated once.

#### Petiole inoculation and sampling

The petioles of cvs Darmor-*bzh* and Bristol were inoculated as described by Dutreux *et al*. [49]. Plants were grown in individual pots and the petioles of the second and third true leaves were inoculated with 8 μL of 10^7^ spores.mL^−1^ pycnidiospore suspensions. Petioles were sampled at 0 (mock), 7 and 14 DPI. At each date, the two inoculated petioles of four plants were cut at their base and placed in Petri dishes. Each petiole was cut 1 cm away from the point of inoculation, and the eight upper fragments obtained were pooled in a 15 mL Falcon tube. The contents of this tube constituted one sample of the upper part of the petiole. All samples were immediately frozen in liquid nitrogen and stored at −80°C until extraction. Two independent replicates were recovered at each time point and the whole experiment was repeated.

#### Petiole inoculation and stem sampling

Plants from the three varieties were sown in individual pots and grown for three weeks before inoculation (S6 Text). When the plants reached the three-leaf stage, the petiole of the second leaf was cut 0.5 cm from the stem. We applied 10 μL of pycnidiospores (10^7^ spores mL^−1^) or sterile water (for mock inoculation) to the petiole section. The plants were kept in the dark, under high humidity, for 36 h, and were then transferred to a growth chamber (S6 Text). Three biological replicates were performed. Each replicate consisted of three blocks of five plants per variety for RNA-Seq sampling. At 14 and 28 DPI, a 0.5 cm-long section of stem was excised from each plant with a razor blade. The stem section was harvested at the level at which the inoculated petiole inserted into the stem, at 14 DPI, and 0.5 cm below this level at 28 DPI. The tissues harvested from each block at each time point (five individual plants) were pooled as a single sample. Harvested tissues were immediately frozen in liquid nitrogen and stored at −80°C.

#### Crop residue sampling

In 2012-2013, the susceptible cultivar Alpaga was sown at Grignon, France, where it was subject to natural infection. Disease incidence, assessed in the fall, revealed high levels of infection, and pre-harvest assessments revealed high levels of disease severity, variable between plants (S6 Text). After harvest, the stem bases of the plants were removed from the field and kept outside until use, either to re-inforce the natural inoculum for the 2013-2014 field assay (see below), or for RNA extraction. For RNA extraction, only plants presenting necrosis on more than 50% of the stem area were used, and stem residues were collected at seven time points after harvest. For one sampling date, two stem residues were pooled to constitute a single sample, and four samples (four replicates) were used per date for RNA sequencing.

#### Stem bases from field sampling

A field experiment was set up at Grignon, France in the fall of 2013, with two cultivars, Darmor-*bzh* and Bristol, with natural, but reinforced inoculum (S6 Text). Stem base tissues (1 cm above the collar of the plants) were collected from both cultivars in the field, at six time points, for RNA extraction (11/20/2013; 12/18/2013; 02/13/2014; 03/13/2014; 04/04/2014; 05/14/2014, 07/08/2014, i.e. one week before harvest (S6 Text)). For each cultivar and time point, we collected three samples (each containing three pooled stem bases) for RNA sequencing. Stem canker severity was assessed by G2 scoring, beginning at the April sampling (S6 Text). Only samples from the susceptible cultivar Bristol collected from March to July were kept for RNA-Seq statistical analyses.

#### Young leaves from field sampling

A field experiment was set up at Grignon, France in the fall of 2017. It included three plots of cv Darmor, subjected to natural, but reinforced inoculum (S6 Text). When typical leaf lesions due to *L. maculans* were visible on more than 60% of the plants (16/11/2017), leaves with leaf lesions were harvested. Five disks of leaf tissue with typical leaf lesions were pooled in a 15 mL Falcon tube (S6 Text). All samples were immediately frozen in liquid nitrogen and stored at −80°C until extraction. Pooled disks from one plot constitutes one replicate, and a similar sampling was performed two weeks later.

#### Infection of cotyledons with ascospores

Stem residues bearing mature pseudothecia were placed on a metallic grid over seven-day-old plants (S6 Text). The stem residues were left over the plants for 24 hours, in the dark, under saturation, to promote ascospore ejection. The plants were then transferred to a growth chamber. After 24 h and 48 h, six cotyledons per pot were randomly cut, pooled in a single 15 mL tube, frozen in liquid nitrogen and stored at −80°C. Cotyledons from four selected neighboring pots (i.e., a total of 24 cotyledons; S6 Text) were pooled to produce a single sample of the early stage of cotyledon infection with ascospores. Two samples were obtained for each time point and the experiment was repeated once.

### RNA extraction and sequencing

Cotyledons, leaf disks, and freeze-dried cultures of fungal spores or mycelia were ground with a Retsch MM300 mixer mill in Eppendorf tubes, with one tungsten carbide bead per tube, for 45 s, at 30 oscillations per second. Petioles were ground with a pestle and mortar. Stem bases from the field and stem residues were ground with a Retsch MM300 mixer mill, with zirconium oxide jars and beads, for 40 s, at 30 oscillations per second. Inoculated stem samples were ground in liquid nitrogen, in the presence of steel marbles, with a TissueLyser II (Qiagen).

RNA was extracted with Trizol reagent (Invitrogen, Cergy Pontoise, France), as previously described [18]. RNA-Seq libraries were prepared as previously described [49]. Each library was sequenced with 101 bp paired-end read chemistry, on a HiSeq2000 Illumina sequencer.

### Annotation of genomic data

#### Comparisons of gene annotations between the two published version of the *L. maculans* strain JN3 genome

The reference genome assembly of *L. maculans* isolate v23.1.3 (aka. JN3) was generated in 2007, by Sanger sequencing [26]. Dutreux *et al.* [49] subsequently generated an improved genome assembly, to achieve assembly at the chromosome scale. We compared this new version of the genome sequence with that published in 2011, using BLAT suite tools [77] (S4 Text).

#### Functional annotations

Functional annotations were predicted with BLAST2GO [78]. Genes encoding CAZymes were annotated manually, together with *de novo* prediction with dbCAN tools v3 [79]. PFam domains were annotated with PFamScan software version 1.6 [80]. For discrimination between proteins with and without annotated functions, the BLAST2GO results and PFam annotations were used, but with the terms “hypothetical protein”, “predicted protein” and “uncharacterized protein” discarded. PKS and NRPS genes were predicted with the SMURF online tool, using default parameters [81].

#### Annotation of genes encoding SSPs

We generated a new repertoire of genes encoding SSPs, by first creating a secretome of predicted secreted proteins with three filters. The predicted secretome contained all the proteins with no more than one transmembrane domain predicted by TMHMM version 2.0. [82], and either a signal peptide predicted by SignalP version 4.1 [83], or an extracellular localization predicted by TargetP version 1.1 [84]. The final SSP repertoire was created by applying a size cutoff of 300 amino acids to the predicted secretome. In parallel, EffectorP version 1.0 [85] was applied to the whole predicted secretome and detected four proteins of more than 300 amino acids in length, which were added to the SSP repertoire.

#### Detection of AT-rich regions

AT-rich and GC-equilibrated regions were annotated with OcculterCut [86] and post-processed to avoid the detection of regions that were too small. OcculterCut uses a predefined window of 1 kb to analyze AT/GC content, and this can lead to the detection of small GC-rich-regions within AT-rich regions in the case of isolated genes. We overcame this problem by defining reliable detected regions (10 kb for AT and 6 kb for GC), which we used as anchors for the merging of regions (S6 Text).

### Analyses on genomic data

#### Genomic localization and enrichment in epigenetic marks

For genes in the eight clusters, we performed Chi^2^ tests to detect enrichment in the associated epigenetic marks (H3K4me2, H3K9me3 or H3K29me3), recovered from Soyer *et al*. [38], to compare the distribution of marks in the whole set of genes. Enrichments were considered significant for *p* values < 0.05; all analyses were performed in R.

#### Gene Ontology enrichment analysis

The Bingo plugin of Cytoscape software v3.5.1 [87] was used for Gene Ontology (GO) enrichment analysis. For each gene, the GO identifier was generated with BLAST2GO software. Significant enrichments in “Molecular Function”, “Biological Process” and “Cellular Component” were detected with a hypergeometric test, by comparing the proportion of genes identified for a given GO function in the subset of interest to that for the whole set of protein-coding genes in *L. maculans*.

#### BLAST

A BLAST alignment was generated with non-redundant protein sequences from the GenPept, Swissprot, PIR, PDF, PDB, and NCBI RefSeq databases. Only the 10 best BLAST hits were selected, with the max_target_seqs: 10 and max_hsps: 10 options.

### Analyses on transcriptomic data

#### Mapping

Raw RNA-Seq reads were mapped with STAR software version 020201 [88]. As *Leptosphaeria biglobosa* isolates could be present in the field samples, we assessed the mapping specificity of the final parameters for the genome of *L. maculans* (S1 Text, **S1 Table**). We chose a number of two base-pair mismatches for a one-step process of mapping onto the concatenated genome of the two species for the final analysis. Based on the maximum intron size within the genome, we allowed for an intron size of 10 000 bp. The other parameters used were as follows: outFilterMultimapNmax: 100; SeedSearchStartLmax: 12; alignSJoverhangMin: 15; alignIntronMin: 10. We then selected the correctly paired reads in BAM files with Samtools v1.6 [89]. Finally, FeaturesCounts version v1.5.1 [90] was used to quantify gene expression for the uniquely mapped and paired reads.

#### Sample correlation analysis

Pairwise Pearson’s correlation analysis was performed on all samples, with the Log2(FPKM +1)-transformed expression values for the 32 selected sets of conditions. The resulting correlation matrix was clustered with the Ward.D2 method. PCA was performed with the FactoRmineR R package [91] on the scaled Log2(FPKM +1) expression values for the four groups identified in the correlation matrix.

#### Detection of differentially expressed genes

Differential expression analyses were performed with the EdgeR package v3.20.9 [92]. Genes with > 30 reads in at least one condition were kept for statistical analysis. Samples were normalized by the TMM method. We fitted a negative binomial generalized linear model to the data with the glmFIt function. We then used the glmLRT function to compare gene expression in each of the 22 *in planta* conditions with that in the 10 *in vitro* conditions, using appropriate coefficients and contrasts (−1/10 for the 10 *in vitro* conditions and +1 for the tested conditions). We also used coefficients to mask the effects of replicate on cotyledon, petiole and stem samples from controlled conditions. Only genes overexpressed relative to the 10 *in vitro* conditions with LogFC > 4 and a *p*-value < 0.01 were selected.

#### Clustering

For the clustering step, genes with a FPKM > 2 in at least one condition were Log2(FPKM +1)-transformed and scaled. The whole 1207 gene set overexpressed in at least one set of conditions *in planta* was extracted from this transformed count table and the self-organizing map method of the Kohonen R package v3.0.8 [93] was used to classify those genes into eight clusters, with 200 iterations.

#### Detection of co-expressed genes

For each cluster, the mean scaled Log2(FPKM +1) value for SSP gene expression was calculated and used as a reference for an analysis of linear regression against the scaled Log2(FPKM +1) expression of all genes. Genes with an R-squared value > 0.80 and an adjusted *p-*value < 0.05 were considered to be highly correlated.

## Acknowledgments

We would like to thank Christophe Montagnier and the experimental unit team, INRAE, France, for the setting up and management of the experimental rapeseed fields; Guillem Rigaill from the Institute of Plant Sciences Paris-Saclay, France, for helpful comments and advice on statistics, and Antoine Gravot, (INRAE IGEPP, France), for helpful discussions and review of the manuscript. For preliminary analyses and genomic work, we would like to thank Fabien Dutreux (CEA, Genoscope, Evry, France). This work was supported in part by Promosol (METAPHOR project), and France Génomique (Leptolife project). The “Effectors and Pathogenesis of *L. maculans*” group benefits from the support of Saclay Plant Sciences-SPS (ANR-17-EUR-0007).

## Author contribution

Conceptualization, RD, TR, MHB; Data curation, EJG, NL, CDA, JMA, CC; Formal analysis EJG, JLS, NL, CDA, JMA; Funding acquisition, PW, TR, MHB; investigation, EJG, NL, JL, CC, AL, JL, MHB; methodology, EJG, JL; Project administration, TR; resources, EJG, NL, CDA, PW, JMA, CC, MHB; supervision, PW, JMA, TR, MHB; validation, JLS, IF; TR; visualization; EJG, JLS; writing draft, EJG, JLS, NL, TR, MHB; writing review, EJG, JLS, NL, IF, CDA, PW, JMA, CC, RD, TR, MHB.

