## Supplemental Figure 1C for "Large-scale transcriptomics to dissect two years of the life of a fungal phytopathogen interacting with its host plant"

**C**

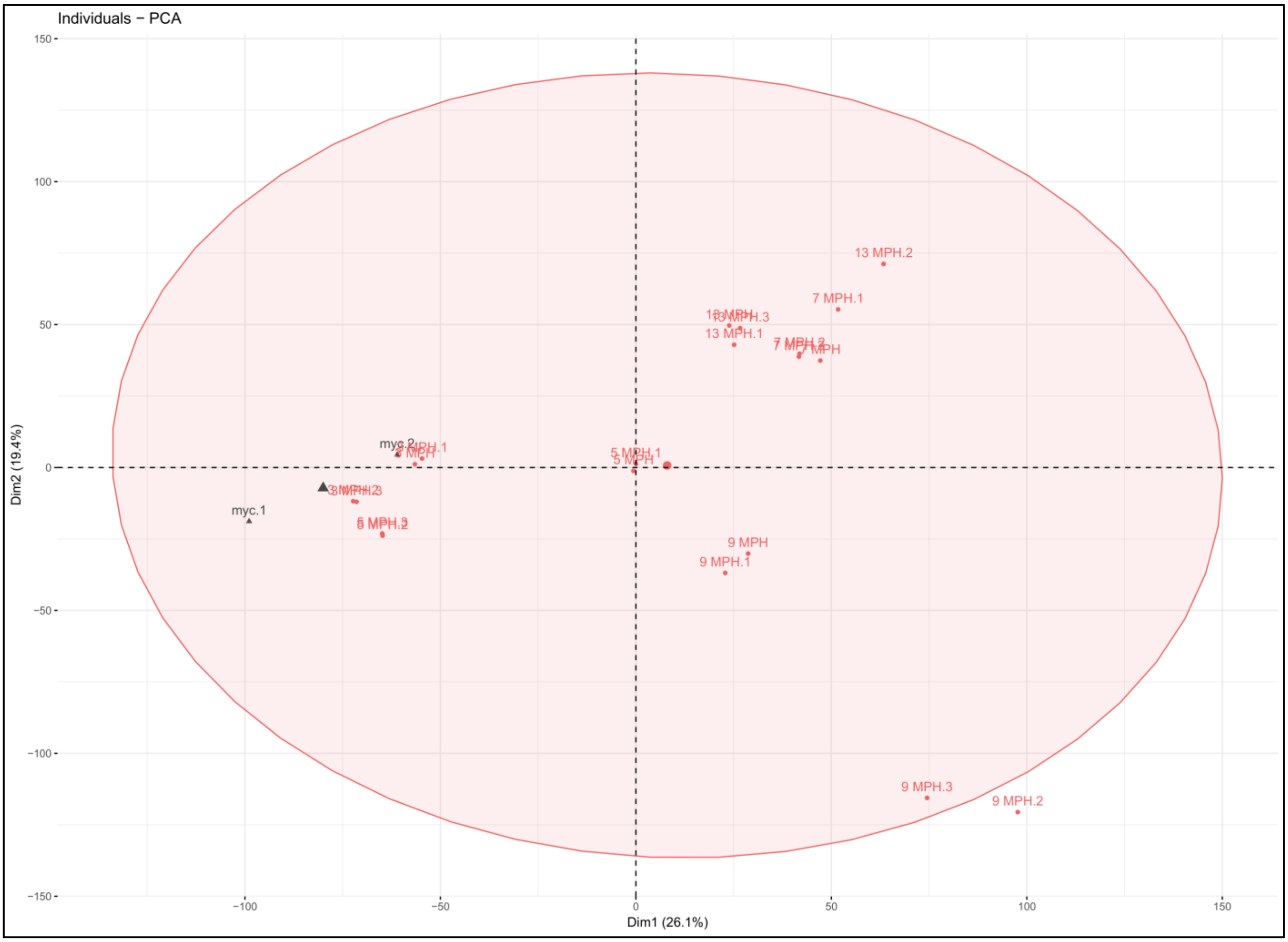


Groups

- IV mycelium (solid growth)
- RF residues

**S1 Fig. Principal component analyses (PCA) of RNA-Seq replicates.** Genes with an FPKM count >2 in at least two sets of conditions were selected and the $Log_2$(FPKM+1) value of each replicate was used as an input for PCA. PCA was performed separately for each group (GP1 to GP4) identified in the correlation analysis (Fig 2), and the results are shown in (A) to (D). Samples are named according to Fig 1 and the last number indicates the replicate number. The two axes represent the first and second principal components. In each PCA, a color is attributed to each independent set of *in vitro* or *in planta* time-course experiment. The ellipses represent the variability of each group, with a confidence interval of 95%.
