## Supplemental Figure 2 for "Large-scale transcriptomics to dissect two years of the life of a fungal phytopathogen interacting with its host plant"

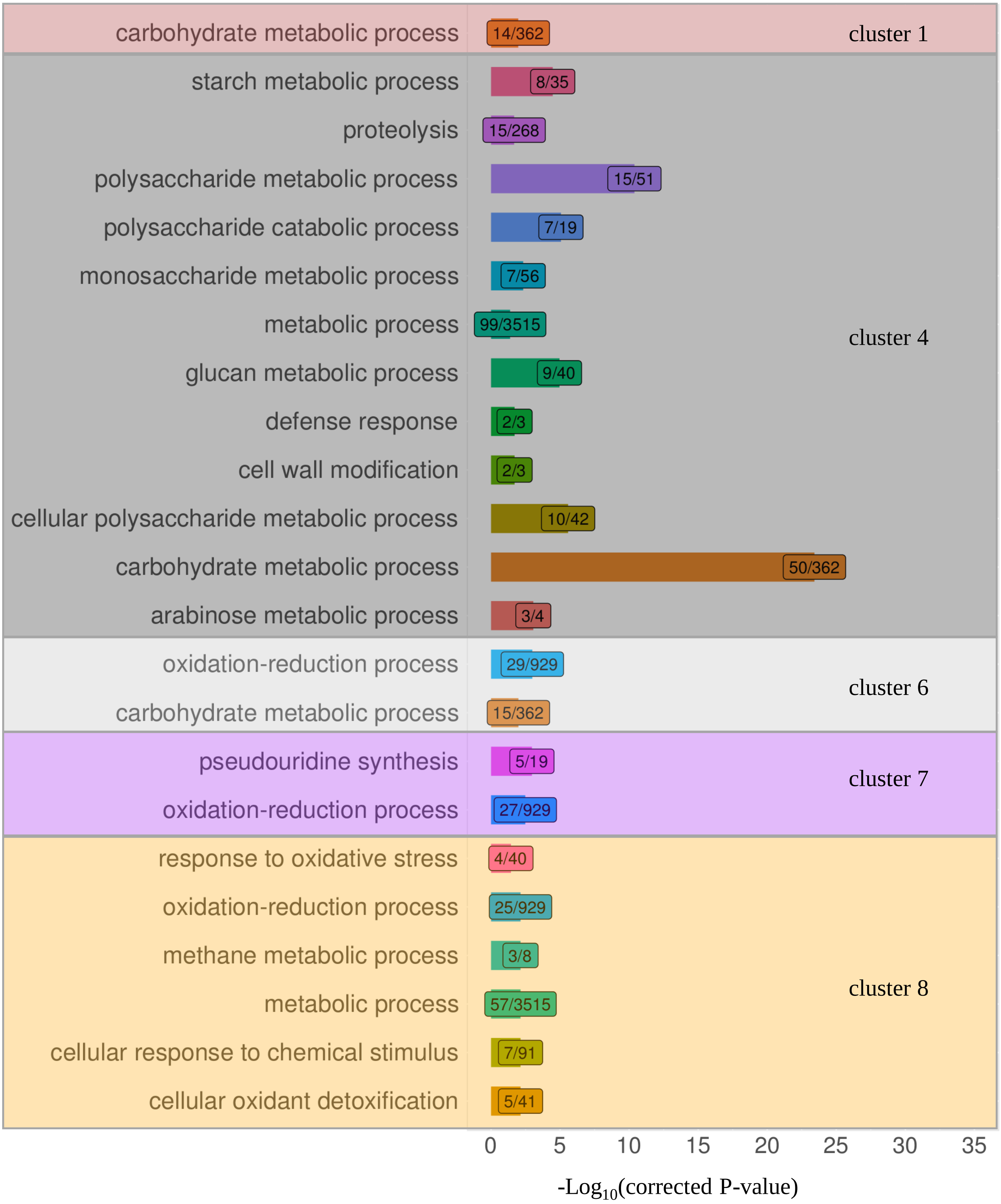


**S2 Fig. Detection of Gene Ontology enrichment ("Biological Process" category) in each of the eight clusters containing the 1,207 *Leptosphaeria maculans* genes overexpressed in at least one set of conditions *in planta* relative to the 10 sets of *in vitro* conditions.** For each cluster, enrichment in a Biological Process category was assessed in a hypergeometric test, with the Cytoscape tool Bingo [87]. The *y*-axis indicates the overrepresented Biological Process terms. The *x*-axis represents the resulting -$Log_{10}$(corrected p-value) of the enrichment test. The numbers in the boxes indicate the number of genes assigned to the corresponding Biological Processes in the cluster (left) and the total number of genes associated to this Biological Process for the whole gene set (right). No significant enrichment in Biological Processes was found in clusters 2, 3 and 5.
