## Supplemental Figure 3 for "Large-scale transcriptomics to dissect two years of the life of a fungal phytopathogen interacting with its host plant"

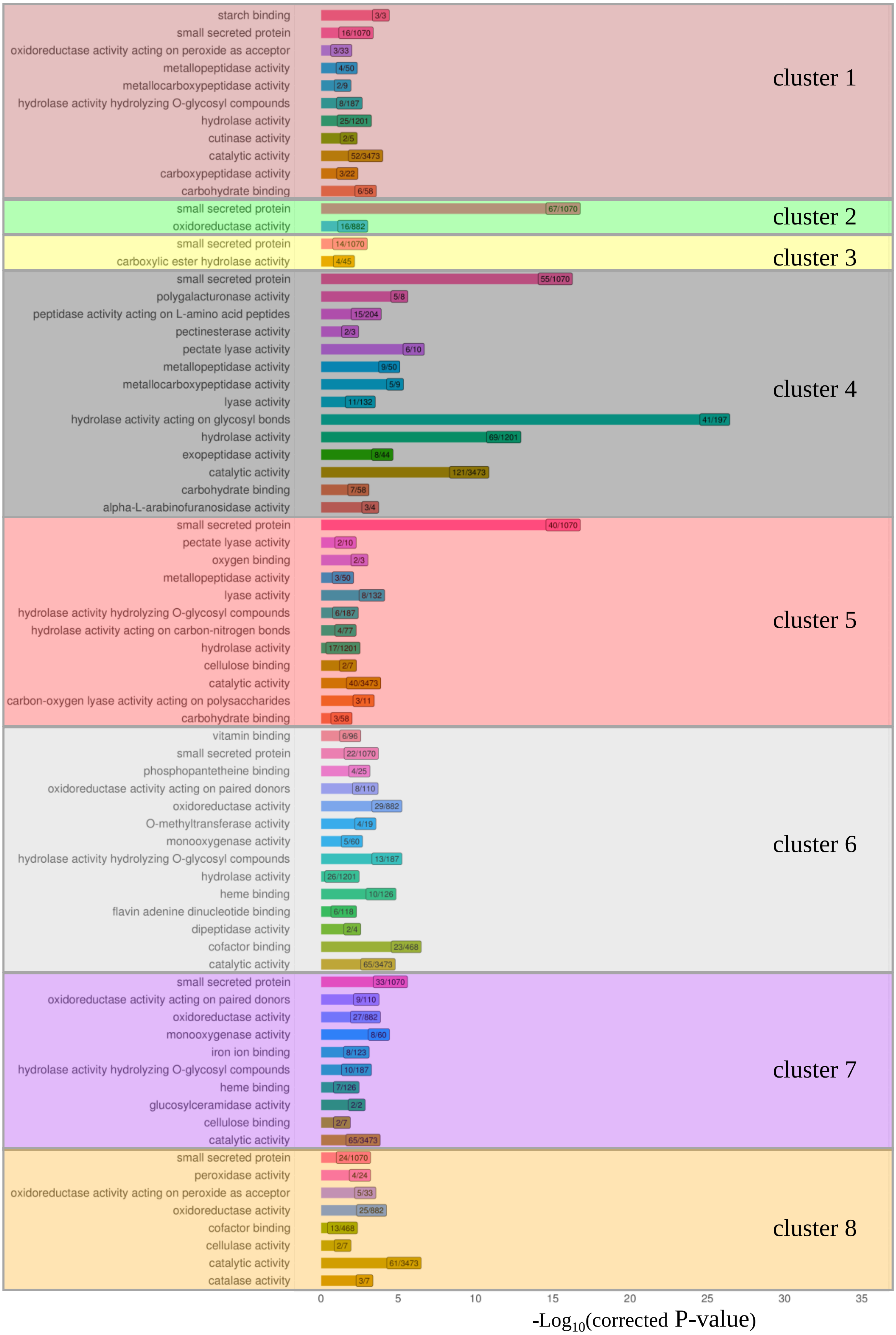


**S3 Fig. Detection of Gene Ontology enrichment ("Molecular Function" category) in each of the eight clusters containing the 1,207 *Leptosphaeria maculans* genes overexpressed in at least one set of *in planta* conditions relative to the 10 sets of *in vitro* conditions.** For each cluster, enrichment in a particular Molecular Function category was assessed in a hypergeometric test, with the Cytoscape tool Bingo [87]. The *y*-axis indicates the overrepresented Molecular Function terms. The *x*-axis represents the -Log$_{10}$(corrected *p*-value) of enrichment for each Molecular Function. The numbers in the boxes indicate the number of genes assigned to the corresponding Molecular Function in the clusters (left) and the total number of genes associated to this Molecular Function term in the whole gene set (right). The 'small secreted protein' category, which is not present in the GO database, was added to the statistical tests.
