## Supplemental Figure 4 for "Large-scale transcriptomics to dissect two years of the life of a fungal phytopathogen interacting with its host plant"

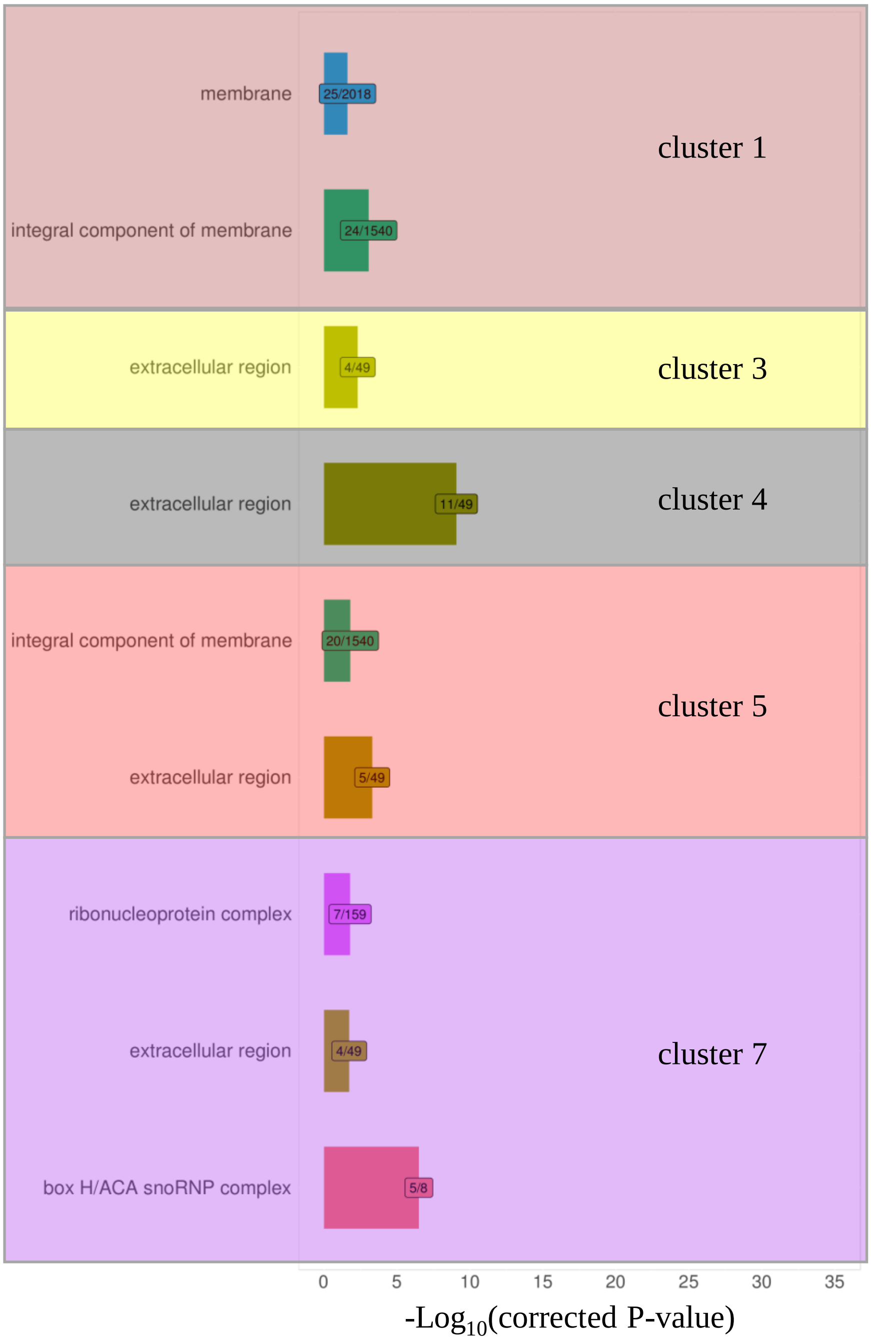


**S4 Fig. Detection of Gene Ontology enrichment ("Cellular Component" category) in each of the eight clusters containing the 1,207 *Leptosphaeria maculans* genes overexpressed in at least one set of conditions *in planta* relative to the 10 sets of conditions *in vitro*.** For each cluster, we assessed the enrichment in a particular Cellular Component category in a hypergeometric test with the Cytoscape tool Bingo [87]. The *y*-axis indicates the overrepresented Cellular Component terms. The *x*-axis represents the -$\text{Log}_{10}$(corrected *p*-value) of the enrichment for each corresponding Cellular Component. The numbers in the boxes indicate the number of genes assigned to the corresponding Cellular Component in the cluster (left) and the total number of genes associated to this Cellular Component term in the whole gene set (right). No significant enrichment in Cellular Components was found in clusters 2, 6 and 8.
