## Supplemental Figure 5 for "Large-scale transcriptomics to dissect two years of the life of a fungal phytopathogen interacting with its host plant"

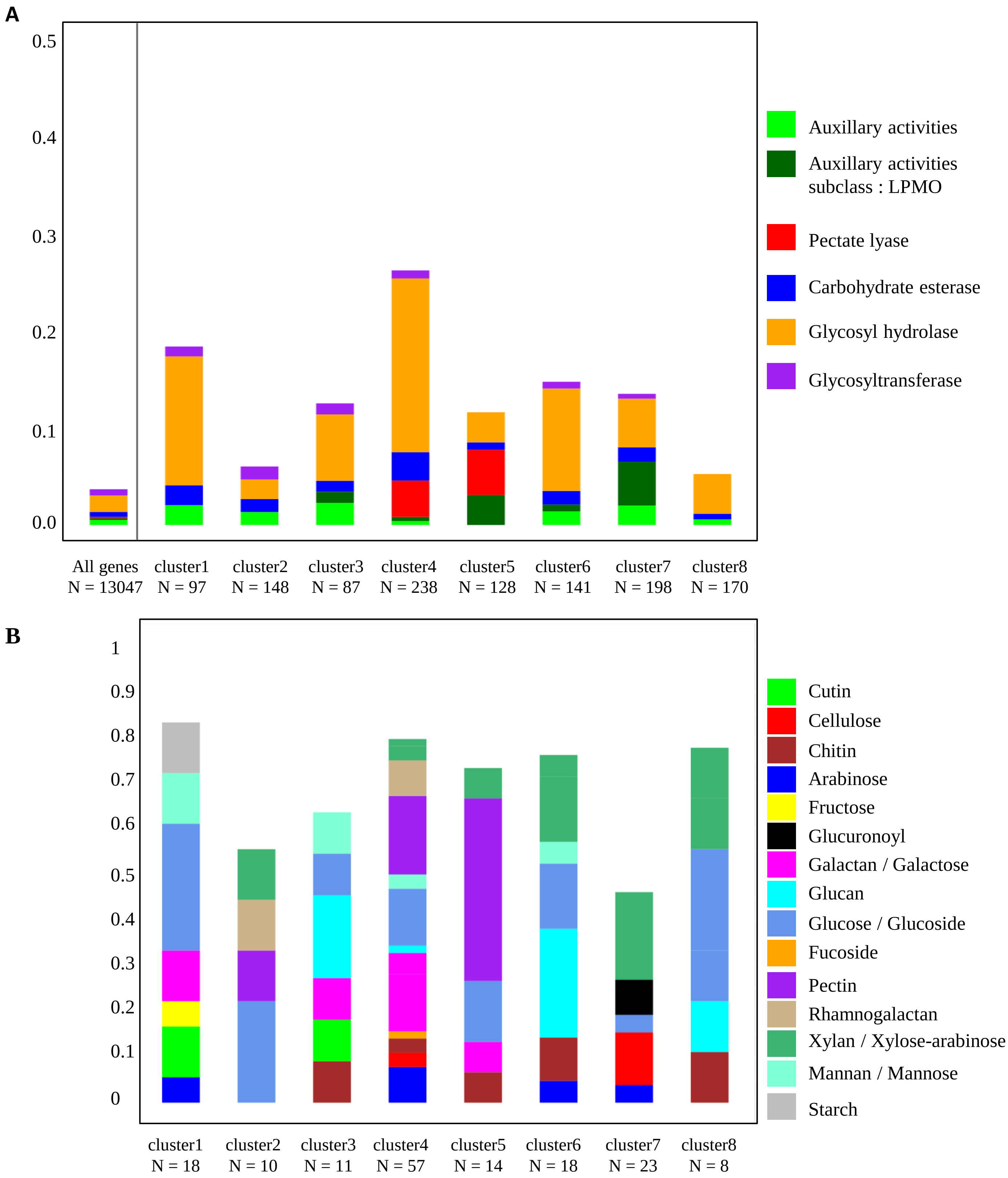


**S5 Fig. Proportion of different classes of CAZymes and their substrates in the eight gene clusters.** (A) The proportion of the five CAZyme classes and one subclass of CAZymes with auxiliary activities (LPMO: lytic polysaccharide monooxygenase) within the clusters (cluster 1 to 8) or the whole gene set of *Leptosphaeria maculans* (*N*, number of genes in each cluster).

(B) The proportion (*y*-axis: from 0 to 1) of the identified CAZyme substrates relative to the total number of CAZymes in each cluster (N).
