## Supplemental Figure 6 for "Large-scale transcriptomics to dissect two years of the life of a fungal phytopathogen interacting with its host plant"

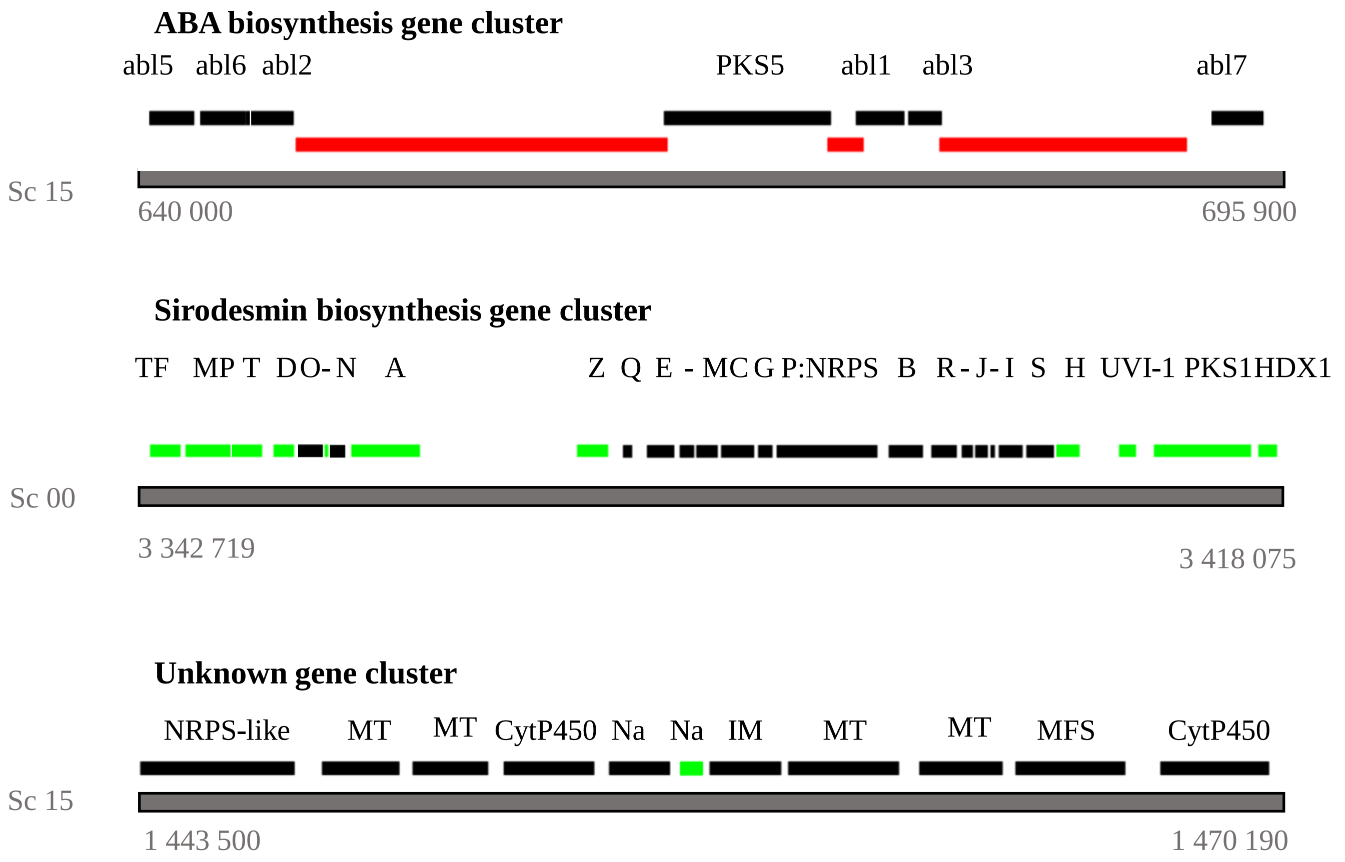


**S6 Fig. Details of the three *Leptosphaeria maculans* secondary metabolite biosynthesis gene clusters upregulated during rapeseed infection.** The genomic regions and scaffolds are represented by the gray bars, with the coordinates indicated beneath them. The red bars correspond to AT-rich regions. The annotations of the genes involved in the abscisic acid (ABA) or sirodesmin biosynthesis are indicated. One secondary metabolite (SM) biosynthesis gene cluster was annotated *de novo* and its borders were delimited by the co-expression of 10 neighboring genes, with one gene encoding an NRPS-like protein. The genes in green represent the genes annotated in the SM cluster in previous studies but not found to be overexpressed during rapeseed infection. The genes in black represent the genes found to be overexpressed during rapeseed infection and present in the cluster. The gene names are indicated for the ABA [27] and sirodesmin biosynthesis cluster [94]. For the *de novo* annotated cluster, the functional annotations predicted by BLAST2GO are indicated (MT: methyltransferase, IM: integral membrane, TF: transcription factor, MP: maltose permease, CytP450: cytochrome P450, Na: no predicted function, MFS, major facilitator superfamily).
