## Supplemental Figure 7 for "Large-scale transcriptomics to dissect two years of the life of a fungal phytopathogen interacting with its host plant"

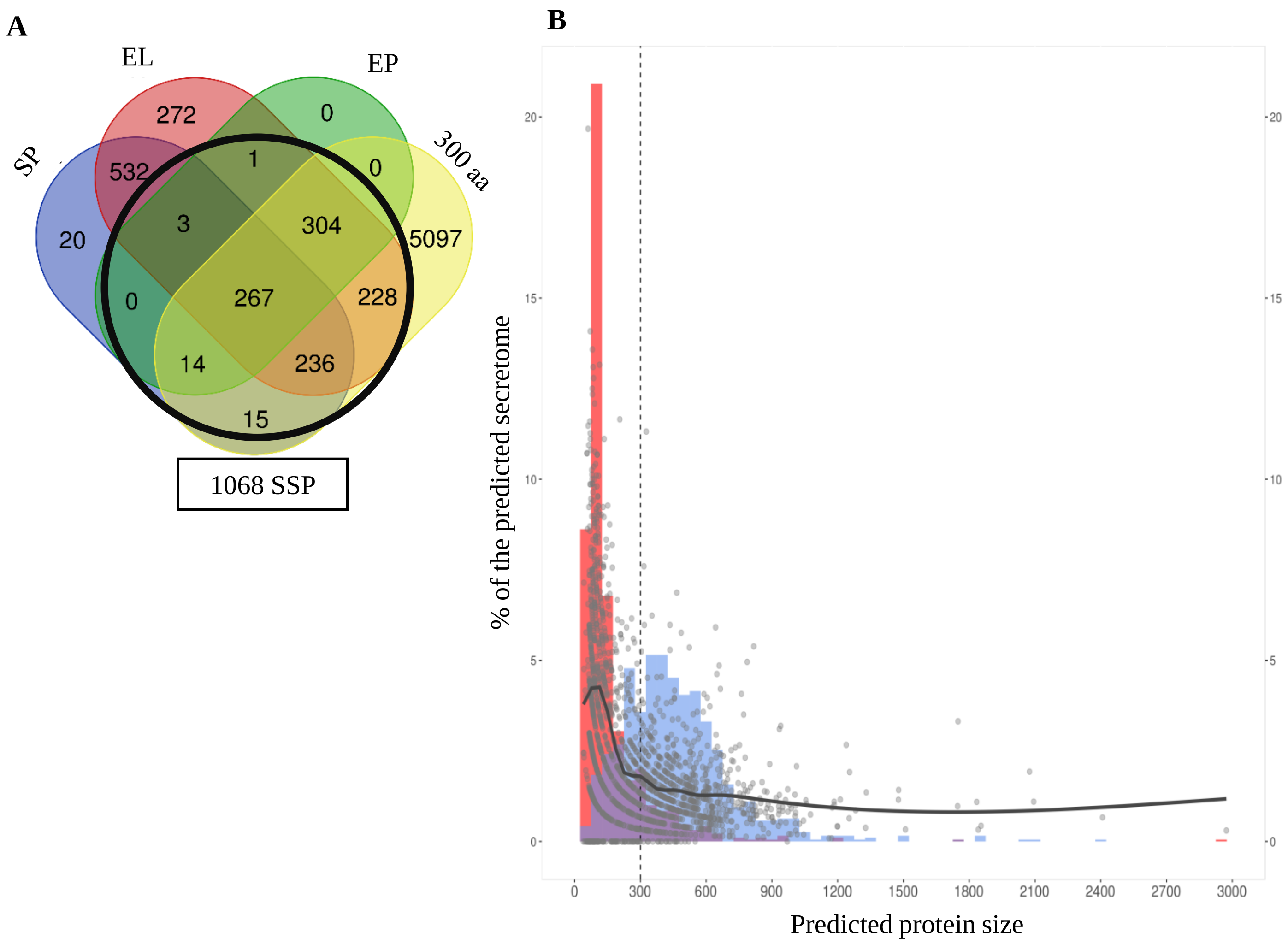


**S7 Fig. Characteristics of the small secreted proteins (SSP) predicted from the *Leptosphaeria maculans* protein set.** (A) Protein features used to predict the SSP repertoire. The Venn diagram contains all proteins with no more than one transmembrane domain. The numbers of proteins with a predicted signal peptide (SP, predicted by SignalP), a predicted extracellular localization (EL: predicted by TargetP), a size of less than 300 amino acids (300 aa), and predicted to be SSPs by the EffectorP tool (EP) are indicated. The black circle includes all proteins comprising the SSP repertoire. (B) Relationship between functional annotation, percentage of cysteine residues and protein size of the 1 892 proteins of the *L. maculans* secretome. The predicted secretome includes all proteins with no more than one transmembrane domain, with a predicted signal peptide or a predicted extracellular localization. The percentage of cysteine residues was calculated (gray dot) and the trend line obtained with the GAM function (generalized additive models with integrated smoothness estimation) is plotted to indicate the change of the percent cysteine residues with protein size. The histogram represents the percentage of genes with and without a predicted function, in blue and red, respectively. The dotted line represents the protein size cutoff (300 amino acids) applied to the secretome.
