## Supplemental Figure 8 for "Large-scale transcriptomics to dissect two years of the life of a fungal phytopathogen interacting with its host plant"

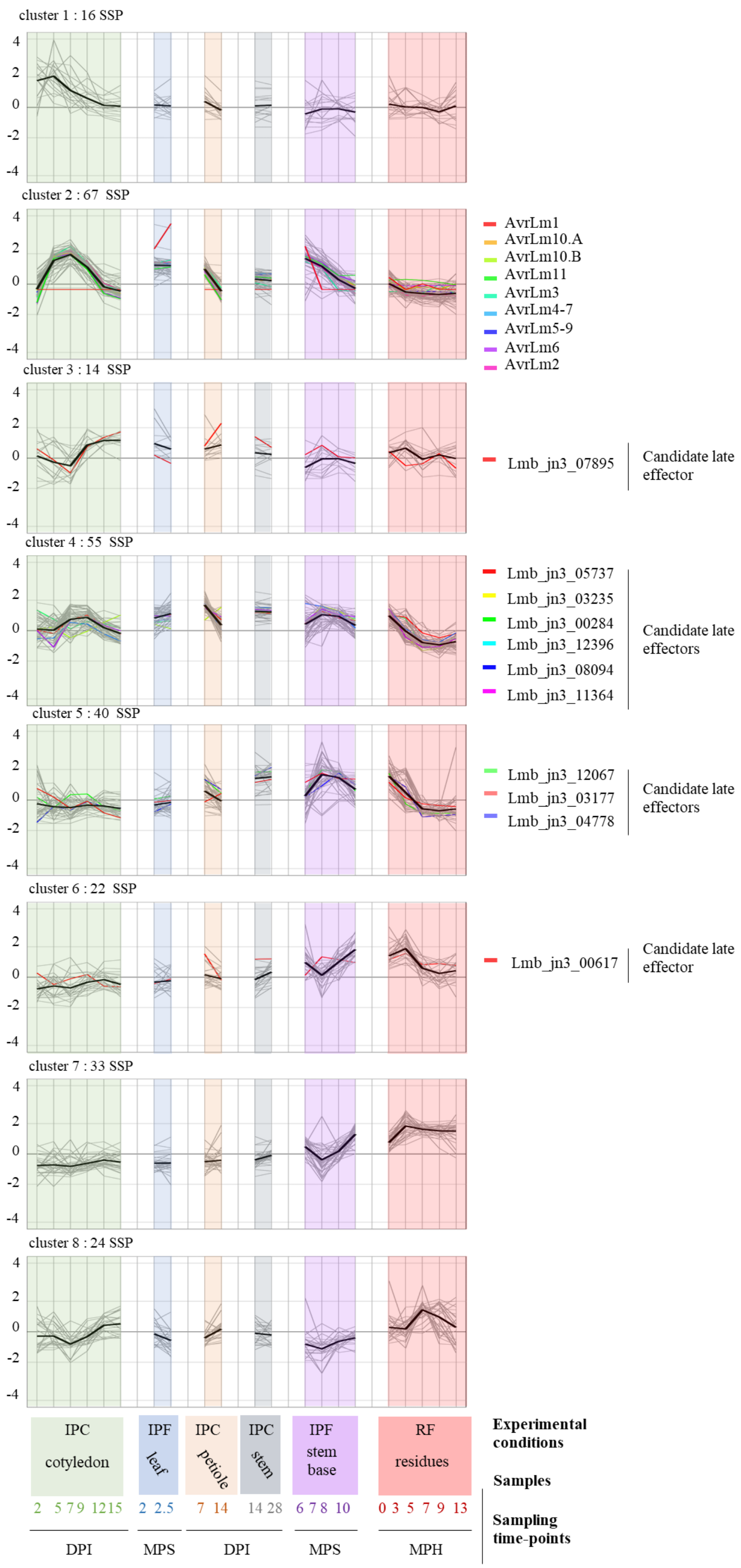


**S8 Fig. Expression of the 271 small secreted protein-encoding genes upregulated in at least one of the 22 sets of conditions *in planta*.** The scaled $Log_2$(FPKM+1) expression values for the 271 SSP genes upregulated in at least one set of conditions *in planta* condition are shown, grouped according to cluster assignment. The distribution and expression of the nine avirulence effector genes (*AvrLm*), and the eight late candidate effector genes [14] are highlighted. The mean expression level of the SSP genes in each of the eight clusters is plotted (black bold curve) and the total number of SSP genes in each cluster is indicated. Three sample features are described: (i) the experimental conditions: IPF, *in planta* field conditions; IPC, *in planta* controlled conditions; RF, residues in field conditions, (ii) the type of plant tissue sampled and (iii) the sampling time points (DPI, days post-inoculation; MPS, months post-sowing; MPH, months post harvest).
