## Supplemental Text 1 for "Large-scale transcriptomics to dissect two years of the life of a fungal phytopathogen interacting with its host plant"

**S1 Text. Characteristics of biological and RNA samples.**

The RNA sequencing dataset represents a highly heterogeneous series of RNA samples that confronted us with two challenges: (i) discrimination of *L. maculans* reads from those of the plant and, mostly, those of other fungal species present at the same time during infection of the plants in agricultural settings, of which the most challenging was *Leptosphaeria biglobosa*, a closely related species to *L. maculans* always associated with it in the wild, (ii) variable amounts of *L. maculans* reads in the samples that may preclude statistical analyses.

(i) Because *L. maculans* belongs to a species complex together with *L. biglobosa*, the field-infected samples, or cotyledons infected with ascospores ejected from "wild" pseudothecia differentiated on residues from the field, are expected to contain both species. *L. biglobosa* was used as a template to design a mapping protocol minimizing wrong assignations and mapping biases (S1 Table). Eventually, our choice was to align reads of all RNAseq samples on the two concatenated genomes in a one-step mapping with two mismatches allowed. These parameters allowed us to reduce the percentage of reads of *L. maculans* wrongly assigned to *L. biglobosa* to 0.008 % of the reads (S1 Table).

(ii) The amount of reads obtained per sample ranged between 16 M and 137 M, with RNAseq samples corresponding to axenic growth (i.e., only consisting of *L. maculans* reads) ranging between 16 M and 25 M reads, and those corresponding to interaction with the plant ranging between 41 M and 137 M reads (S2 Table). The number of reads assigned to *L. maculans* in the samples was very variable, and in some cases insufficient to be submitted to statistical analyses. This mainly regarded infection of cotyledons with ascospores from the field, with less than 0.01% of the reads that were assigned to *L. maculans*, and the earliest samplings of stem bases of plants naturally infected in the field (November to February samplings, two to five months post-sowing - MPS) in which the quantity of reads assigned to *L. maculans* was very low and ranged from 1153 reads (0.002%) to 23 934 reads (0.028 %) (S2 Table). These samples were discarded for the statistical analyses, but they showed that the stem base colonization begins early in the plant development, and that the fungus shows a steady increase of transcriptomic activity bursting at the end of winter. The active presence of *L. maculans* on residues leftover for more than one year was obvious on all residues, but, again the number of reads attributed to *L. maculans* was highly variable, roughly ranging between 0.18% and 58% of the sample libraries. On an inert substratum, devoid of living plant material (and RNA reads from the plant) this confirms the presence of a dynamic community of Eukaryote microbes [55]. In conclusion, 87 samples showed a percentage of reads assigned to *L. maculans* ranging between 0.08% and 87.01% and were useable for statistical analyses. These corresponded to 32 different conditions.
