## Supplemental Text 2 for "Large-scale transcriptomics to dissect two years of the life of a fungal phytopathogen interacting with its host plant"

**S2 Text. Reproducibility between replicates.**

Variability between replicates was analysed with a PCA approach and indicated consistency between replicates in most of the samples (S1 Fig). The highest variability was detected between replicates from asymptomatic field-infected stem bases (S1A Fig) or crop residues (S1C Fig) probably linked to numerous environmental factors causing transcriptomic variability from one replicate to another or variable intensity of fungal colonization from one plant to the other. One series of *in vitro* growth conditions, those promoting differentiation of pycnidia and/or pseudothecia, also showed some variability between replicates (S1D Fig) that may reflect lack of synchronicity from one sample to the other in the differentiation processes.
