## Supplemental Text 3 for "Large-scale transcriptomics to dissect two years of the life of a fungal phytopathogen interacting with its host plant"

**S3 Text. Generation of a new repertoire of small secreted proteins considered as candidate effectors.**

The secretome of *L. maculans*, defined as proteins with (i) at most one transmembrane domain (predicted by TMHMM) and (ii) a predicted signal peptide (predicted by SignalP) or a predicted extracellular localization (predicted by TargetP) encompassed 1,984 predicted proteins in the v2 genome and annotation recently published by Dutreux *et al*. [49]. Along with abundance of cysteine residues, and scarce homologies in databases, a size cut-off is often applied to refine the secretome in candidate effectors, but the relevance of this criterium has sometimes been questioned in the literature. Here, we applied a size cut-off of 300 amino-acid, resulting in a set 1,064 small secreted proteins (SSP) (S7A Fig).

To question and assess the impact of this size cut-off, we analysed the percentages of cysteine residues and of predicted function in the whole predicted secretome depending on protein size (S7B Fig). The cut-off at 300 amino-acid discarded 73 % of genes with a functional annotation and kept a set of proteins with a significant enrichment in cysteine residues compared to the whole secretome.

EffectorP, used on the whole secretome, predicted that 596 proteins of the secretome are effector candidates of which 592 are among the 1,064 SSP identified with the 300 AA size cut-off and only four are larger than 300 AA (S7A Fig). These four proteins were added to the 1,064 to generate the new repertoire containing containing 1068 proteins (S7B Fig). This set contained all known *AvrLm* genes, but two of the eleven late effectors analysed by Gervais *et al.* [14] were discarded because they did not match all the criteria and they were manually added to the SSP set leading to a final repertoire of 1,070 genes. The results of each tool used to create the SSP repertoire as well as other features such as the corresponding gene annotation in v1 annotation [26], the genomic localization, the functional annotation, the number of cysteine residues and the *AvrLm* gene Id. are detailed in S5 Table. This repertoire is enriched in genes located in AT-rich regions (S6 Table).
