## Supplemental Text 4 for "Large-scale transcriptomics to dissect two years of the life of a fungal phytopathogen interacting with its host plant"

### S4 Text. Comparison between the v1 and v2 versions of gene model annotations.

Changes in genome assembly and gene annotations between 2011 and 2018 [26,49] showed drastic changes in the gene repertoire: while 88.3% (11,154) of the genes of v1 annotation showed at least one overlap with genes of v2, only 59% had a unique overlap and 29.2 % were merged or splitted in the v2 annotation. In addition 11.7% of gene models of v1 had no correspondence with the v2 annotation while 1,905 genes of v2 were absent from the previous annotation. Due to these changes in gene annotation, we generated a new repertoire of candidate effectors.
