## Supplemental Text 5 for "Large-scale transcriptomics to dissect two years of the life of a fungal phytopathogen interacting with its host plant"

**S5 Text. Genes expressed when ascospores colonize the plant tissues or when the fungus is dormant in the stem.**

Among the 37 conditions from which RNA samples were obtained, five were discarded from statistical analyses due to the low amount of fungal material leading to insufficient number of *L. maculans* reads. They corresponded (i) to ascospore germination and growth on cotyledons after 24 and 48h of putative ejection of the ascospores, and (ii) to oilseed rape stem bases collected in the field 2, 3 and 5 months after sowing, i.e. during the winter period.

We nevertheless tried to take advantage of these data to identify wether these genes could assigned to one or the other wave of expression. For germinating ascospores, only 86 genes showed more than 10 RNAseq reads in one or the other sample (S7 Table) and 51 of them were found in one of the expression clusters. The majority, 32 genes (60.7%), were included within cluster 2, 83.8% of them being SSP genes of cluster 2 (including *AvrLm3*, *AvrLm4-7*, *AvrLm5*, *AvrLm6*, *AvrLm10A*, *AvrLm10B* and *AvrLm11*). Twenty-seven % of the genes were included within clusters 7 or 8. Interestingly, these latter corresponded exclusively to genes detected 24 h after ascospore inoculation while genes of cluster 2 were exclusively found in samples from 48 hpi. Similarly, in samples from stem bases, 41, 50 and 394 genes showed more than 10 RNA-Seq reads in one or the other replicate for November (3 MPS), December (4 MPS) and February (6 MPS), respectively. 104 (26.2%) of these genes were included in the expression clusters (S7B Table), and mostly belonged to cluster 2 (67 genes including 40 genes encoding SSPs, including *AvrLm2*, *AvrLm3*, *AvrLm4-7*, *AvrLm5*, *AvrLm6*, *AvrLm10A*, *AvrLm10B* and *AvrLm11*).
