## Supplemental Text 6 for "Large-scale transcriptomics to dissect two years of the life of a fungal phytopathogen interacting with its host plant"

**S6 Text. Additional material and methods.**

### *In vitro* culture

For the mycelial state on solid medium, the fungus was cultivated on V8-agar plates. A plug of an actively growing culture was placed on the centre of 9 cm diameter V8-agar Petri dishes. The plates were sealed with Parafilm and were incubated at 25°C in the dark. After 7 days of growth the aerial mycelium was collected with a spatula and immediately frozen with liquid nitrogen in an Eppendorf tube. The mycelia from a minimum of three plates were pooled in one sample, and mycelia harvested from two independent experiments were submitted to RNA sequencing.

To produce pycnidiospores, the fungus was cultivated on V8-agar plates without Parafilm and under both white light and near-UV light with a 12 hour photoperiod. After 10 days, spores were collected as previously described [71]. The spore suspension from 9 Petri dishes were pooled in 50 mL Falcon tubes and centrifuged for 15 min at 4°C and 2 600 *g*. The supernatant was discarded and the spores were suspended in 4 mL of sterile water. One mL of this spore suspension was centrifuged similarly in an Eppendorf tube and the pellet was immediately frozen in liquid nitrogen to constitute the "resting spores" sample. For germinating spores, Erlenmeyer containing 20 mL of Fries liquid medium were seeded with the spore suspension at a final concentration of $1.25.10^{8}$ spores.$mL^{-1}$ and submitted to orbital shaking at 150 rotations.$mn^{-1}$. After 24 h at 25°C, the germinating spore suspension was centrifuged in a Falcon tube for 7 min at 9000 *g*, and 4°C, re-suspended in 1 mL of sterile water for a second centrifugation for 5 min at 9000 *g* in an Eppendorf tube. The centrifugate was immediately frozen and stored at -80°C to constitute the "germinating spores" sample.

Mycelium was also obtained in liquid cultures in Fries liquid medium as previously established [102] after seven days of cultivation.

Sexual mating conditions were obtained as previously described [103]. Briefly, plugs of two isolates of the opposite mating types, JN2 and Nz-T4, were deposited side by side on V8 agar plates. After seven days of cultivation under sporulating conditions, 5 mL of 1.5% water agar were poured over the plates, further sealed with Parafilm and placed at 10-11°C under backlight with a 12h photoperiod. Plates with JN2 only were used as a control. Fungal material on the plates was harvested at days 7 (before pouring the agar overlay), 20 and 35 days of cultivation. For each time point, the fungal material (mix of mycelium, pycnidia and maturing pseudothecia) of 10 plates were recovered using a spatula and pooled in one Eppendorf tube, immediately frozen with liquid nitrogen and stored at -80°C.

#### Cotyledon inoculations and sampling

Cotyledons of 10 day-old plants of cv. Darmor-*bzh* were inoculated as described previously [76]. Briefly, 48 seeds were sown per tray (4 rows of 12 plants) and 12-15 days old plantlets were inoculated. Ten µL droplets of $10^7$ conidia.mL$^{-1}$ suspension were deposited on the centre of each half-cotyledon previously punctured with a needle. Plants were maintained in the dark and saturating humidity at room temperature for 48 hours then maintained in growth chambers at 16°C (Night)/24°C (Day) with a 16 hours light photoperiod. Control plants were mock-inoculated with sterile deionized water. Samples for RNA-Seq were recovered at day 0 (with or without wound), 2, 5, 7, 9, 12, 15 and 17 after inoculation. At each time-point, eight cotyledons from eight different plants were randomly selected on the trays. The plant tissues around the inoculation were cut with a 10 mm disposable punch and the 16 corresponding samples were pooled together in a sterile Falcon tube, immediately frozen in liquid nitrogen and stored at -80°C until extraction. At each time point, two replicates were recovered and the whole experiment was repeated once. At each time point and repeat, mock-inoculated samples were recovered similarly.

#### Petiole inoculation and stem sampling

Plants were sown in individual pots (9 * 9* 9.5 cm) in commercial Falienor substrate (65% Irish blond peat, 20% Baltic black peat, 15% perlite, 2% Danish clay) and grown in a growth chamber (16h of light at 18°C at 200 µmol m$^{-2}$ s$^{-1}$ and 8h of dark at 15°C) for three weeks before inoculation. When the plants were at the three-leaf stage, the petiole of the second leaf was cut 0.5 cm from the stem. 10 µL of inoculum (conidia at $10^7$ spores.mL$^{-1}$) or 10 µl of sterile water for mock inoculation were applied on the petiole section. The plants were kept in darkness and high humidity for 36 h and then grown in a growth chamber (16h of light at 20°C at 200 µmol m$^{-2}$ s$^{-1}$ and 8h of dark at 18°C). Sub-irrigation was performed using the commercial fertilizer preparation Liquoplant Bleu 0.3% (Plantin, France).

#### Field experiment establishment, disease assessment and sample preparation

In the 2012-2013 assay, the disease incidence (% of plants with at least one phoma leaf spot) was estimated in November 2012 and reached 80%. Disease severity before harvest was estimated on 50 plants using the “G2” rating score as described previously [75]. Only 4% of the plants displayed no stem canker symptom. For 29%, 40%, 22%, and 5% of the plants, the necrotic area represented less than 25%, between 25 and 50%, between 50 and 75% or more than 75% of the stem section area, respectively, showing variable intensity of plant contamination between plants.

The 2013-2014 field experiment was sown in Grignon, France in autumn 2013 (6 September). The two cultivars Darmor-*Bzh* and Bristol were sown in 10 m x 1.75 m plots in duplicates, at a 60 seeds per $m^2$ density. Naturally infected stem residues of the susceptible cv. Alpaga from the previous cropping season were added over the soil the 19/09/2013 to reinforce the natural inoculum. The disease pressure was measured at two dates in autumn (18/10/2013 and 13/15/11/2013). For that, 2 x 50 consecutive plants per micro plot were observed and each plant was classified in one of the following classes: class 0, no visible phoma leaf spot, class 1, less than 5 leaf spots per plant, class 2, more than 5 leaf spots per plant. At the second rating date, the disease incidence (% of infected plants) reached 92.7% for Bristol and 99% for Darmor-*Bzh*. More than 70% and 90 % of the plants of Bristol and Darmor-*Bzh*, respectively, displayed more than 5 leaf lesions per plant. Whole plants were collected from this field at 6 time points for RNA extraction (20/11/2013; 18/12/2013; 13/02/2014; 13/03/2014; 4/04/2014; 14/05/2014, 8/07/2014 i.e. one week before harvest). For each time point and each cultivar, six plants were collected per micro plot. Plants were washed in the lab with running water and rapidly air dried. Stem section of 1 cm was cut just above the limit between the root and the stem (collar with leaf scares). This piece of stem was then sliced in a sterile Petri dish and the first and last slices were discarded. The central parts of three plants were pooled in one Falcon tube to constitute one sample that was immediately frozen with liquid nitrogen and stored at -80°C before RNA extraction. For one given cultivar and time point, 4 samples were thus collected but only three were submitted to RNA sequencing. The stem canker severity (G2 rating, [75]) was calculated on the collected plants starting from the April sampling, and then on 60 plants per plot the 20$^{th}$ June 2014.

#### Young leaves from field sampling

A field experiment was set up in Grignon, France in autumn 2017. The cultivar Darmor was sawn the 31/08/2017 along with other varieties, in 8 m x1.75 m plots in triplicates, at a 60 seeds per $m^2$ density. In order to reinforce the natural inoculum of the field, naturally infected stem residues of cv. Alpaga from the previous cropping season were added over the soil the 16/09/2017 at a density of 4 residues per $m^2$. The 16/11/2017 typical leaf lesions due to *L.*

*maculans* were visible in more than 60% of the plants. Ten leaves per plot with leaf lesions were harvested. In the lab, leaves were washed with running water, rapidly dried between sterile filter papers. Disks of leaf tissues were cut with a sterilized punch (diameter: 2cm), and classified in three categories: disks without any visible symptoms, disks with atypical symptoms potentially attributable to *L. biglobosa*, and disks with typical leaf lesions. Five leaf discs per category were pooled in a 15 mL Falcon tube. All samples were immediately frozen in liquid nitrogen and stored at -80°C until extraction. Here, pooled disks from typical leaf lesions only were submitted to RNA-Seq.

#### Cotyledon infections with ascospores

Stem residues of the cv. Bristol were collected at the end of the 2013-2014 field experiment (see above) and were left outside from July 2014 to March 2015, allowing pseudothecia to differentiate on the stems. Seven days-old plants of Bristol, grown in 5 x 5 cm pots with 10 plants per pot, were transferred in 50x40 cm plastic bowls (24 pots per bowl). Stem residues were rinsed with water, then allowed to dry up again and placed on a metallic grid over the plants (30 residues per bowl). Stem residues were left 24hours over the plants, maintained in the dark with saturated humidity, to allow ascospores to be ejected on the cotyledons. Then the plants were transferred to growth chambers (18-24°C night-day, 16h photoperiod as above). After 24h and 48h, six cotyledons per pot were randomly cut, pooled in one 15 mL tube, frozen in liquid nitrogen and stored at -80C. The remaining cotyledons were observed for typical leaf lesions one week later. This allowed us to select cotyledons corresponding to pots on which leaf spots, typically due to ascospore due to their rapid development compared to conidia, had developed. Based on this information, cotyledons from 4 neighbour pots on which leaf lesions had developed (i.e., a total of 24 cotyledons) were pooled to produce samples of early stage (24h and 48h) of cotyledon infection with ascospores. Two samples were produced at each time point and the experiment was repeated once.

#### Annotation of isochores

As for the gene models, the new genome version impacted the boundaries of AT/GC isochores previously defined on the first version of the genome assembly [26]. Contrary to the previous annotation, which was manually curated, an automatic protocol was set up combining Occultercut and post-processed to detect coordinates of isochores. Finally, the genome was divided in 553 regions, among them 290 were AT-rich regions (16.01 Mb, 34.8% genome) and 263 were GC-equilibrated regions (29.96 Mb, 65.2% genome).
