## Supplemental Table 1 for "Large-scale transcriptomics to dissect two years of the life of a fungal phytopathogen interacting with its host plant"

**S1 Table. Results of the mapping parameters tested for distinguishing between reads from *L. maculans* and *L. biglobosa.***

[a.] Reads of the two samples from *in vitro* cultures of either *L. maculans* (isolate JN2) or *L. biglobosa* (isolate G12-14) were mapped independently on the genomes of *L. maculans* (isolate JN3) and *L. biglobosa* isolate G12-14)

[b.] Reads of the two samples from *in vitro* cultures of either *L. maculans* (isolate JN2) or *L. biglobosa* (isolate G12-14) were both mapped on one artificial genome created by the concatenation of the *L. maculans* (isolate JN3) and *L. biglobosa* (isolate G12-14) genomes.

| **Reference mapping genome** | **RNAseq samples used** | **Number of mismatch allowed** | **% of reads mapped on *L. maculans* (JN3) genome** | **% of reads mapped on *L. biglobosa* (G12-14) genome** |
|---|---|---|---|---|
| [a.] JN3 or G12-14 genomes | *In vitro* culture of *L. maculans* (JN2) | 0 | 96.62 | 0.46 |
| | | 1 | 97.27 | 0.97 |
| | | 2 | 97.42 | 1.63 |
| | *In vitro* culture of *L. biglobosa* (G12-14) | 0 | 0.76 | 95.13 |
| | | 1 | 1.54 | 95.59 |
| | | 2 | 2.66 | 95.69 |
| [b.] JN3 and G12-14 concatenated genomes | *In vitro* culture of *L. maculans* (JN2) | 0 | 96.7 | 0.04 |
| | | 1 | 97.32 | 0.06 |
| | | 2 | 97.45 | 0.08 |
| | *In vitro* culture of *L. biglobosa* (G12-14) | 0 | 0.10 | 95.15 |
| | | 1 | 0.11 | 95.58 |
| | | 2 | 0.12 | 95.67 |
