## Supplemental Table 3 for "Large-scale transcriptomics to dissect two years of the life of a fungal phytopathogen interacting with its host plant"

**S3 Table. Detection and cluster distribution of PKS and NRPS genes among the 1,207 *Leptosphaeria maculans* genes upregulated *in planta*.**

[a.] PKS and NRPS were annotated by SMURF tool

| Gene ID [49] | Cluster assignation | PKS/NRPS annotation[a] | Associated metabolite |
|---|---|---|---|
| Lmb_jn3_11562 | cluster 2 | PKS | Abscisic acid |
| Lmb_jn3_03490 | cluster 4 | NRPS | - |
| Lmb_jn3_12570 | cluster 5 | PKS | - |
| Lmb_jn3_00988 | cluster 6 | NRPS | Sirodesmin |
| Lmb_jn3_05348 | cluster 6 | PKS | Phomenoic acid |
| Lmb_jn3_06412 | cluster 6 | PKS | - |
| Lmb_jn3_11822 | cluster 6 | NRPS-like | - |
