## Supplemental Table 6 for "Large-scale transcriptomics to dissect two years of the life of a fungal phytopathogen interacting with its host plant"

**S6 Table. Comparison of protein features and genomic localization between the small secreted protein (SSP) repertoire predicted from v2 genome annotation relative to the whole gene set with those obtained for the previously predicted SSP repertoire.**

[a,b] Results of the statistical comparisons made on cysteine percent and proportion of genes localized in AT-rich regions between the whole protein set and the predicted SSP set with a student and a Chi-square test respectively. Asterisks indicate a significant enrichment in cysteine and in gene localized in AT-rich region is detected in the new predicted SSP set. *** : p-value < 0.001, **: p-value < 0.01.

[c,d,e] Results of the statistical comparisons made on CDS size, cysteine percent (with a student test) and on the proportion of genes localized in AT-rich regions (with a Chi-squared test), between previously published [26] and the SSP repertoire predicted in the current study. Asterisks indicate a significant reduction in SSP protein size, a significant increase in cysteine percent, and a significantly lower proportion of genes localized in AT-rich region in the newly predicted SSP set. *** : p-value < 0.001, **: p-value < 0.01, * p-value < 0.05.

| | Predicted SSP on the v2 genome | All proteins | SSP repertoire in [26] |
|---|---|---|---|
| Total number of proteins | 1,070 | 13,047 | 651 |
| Average coding sequence size (AA) | 139.81 | 392.33 | 155.90*** [c] |
| Cysteine composition (%) | 3.51 | 1.98 *** [a] | 3.10** [d] |
| Genes in AT-rich regions (%) | 6 | 1.70 *** [b] | 8.70* [e] |
| Proteins with a predicted function (%) | 25 | 58.58 | 13 |
