## Supplemental Table 7 for "Large-scale transcriptomics to dissect two years of the life of a fungal phytopathogen interacting with its host plant"

**S7 Table. Genes with more than 10 reads in (A) at least one of the four replicates of samples at 24 h or 48 h after ascospore ejection on cotyledons (B) at least one of the nine replicates from the three samples of stem base tissues at 2, 3 or 5 months post sowing.**

[a] Total number of genes  with a count > 10 reads in at least one of the replicates at 24h or 48h post infection by ascospores

[b] No of genes absent from the set of 1,207 genes over-expressed in one of  the 22 analyzable conditions compared to the ten *in vitro* conditions

[c] No of genes present in the set of 1,207 genes over-expressed in one of the 22 analyzable conditions compared to the ten *in vitro* conditions, and their cluster assignation (cluster 1  to cluster 8).

**A**

| **Gene categories** | | **Non-SSP** | **SSP** |
|---|---|---|---|
| Total number of genes with a count > 10 reads [a] |  | 55 | 31 |
| No of genes showing no differential expression [b] | | 33 | 2 |
| No of genes over-expressed during the infectious cycle and their cluster assignations [c] : | | 22 | 29 |
| | cluster 1 | 0 | 1 |
|  | cluster 2 | 6 | 26 (including 7 *AvrLm* genes) |
| | cluster 3 | 0 | 1 |
| | cluster 4 | 2 | 0 |
| | cluster 5 | 0 | 0 |
| | cluster 6 | 0 | 1 |
| | cluster 7 | 5 | 0 |
| | cluster 8 | 9 | 0 |

**B**

| Gene categories | | Non-SSP | SSP |
|---|---|---|---|
| Total number of genes with a count > 10 reads [a] | | 331 | 62 |
| No of genes showing no differential expression [b] | | 278 | 11 |
| No of genes over-expressed during the infectious cycle and their cluster assignations [c] : | | 53 | 51 |
| | cluster 1 | 6 | 2 |
| | cluster 2 | 27 | 40 (including 6 *AvrLm* genes) |
| | cluster 3 | 2 | 2 |
| | cluster 4 | 8 | 3 |
| | cluster 5 | 9 | 4 (including one late effector) |
| | cluster 6 | 0 | 0 |
| | cluster 7 | 1 | 0 |
| | cluster 8 | 0 | 0 |
