## Supplemental Table 8 for "Large-scale transcriptomics to dissect two years of the life of a fungal phytopathogen interacting with its host plant"

**S8 Table. Conservation of the protein sequences encoded by highly co-expressed SSP genes.**

[a.] Number of SSP genes detected as highly correlated with each reference expression wave

[b.] Number of *AvrLm* effectors and *LmSTEE* effectors detected as highly correlated with each reference expression wave

[c.] Number of SSP sequences with no homology after a BLAST search

| Assignation to wave | [a.] Total no of highly co-regulated SSP genes | [b.] No of known effectors in *L. maculans* | [c.] No of SSP without homologous protein in the non-redondant protein NCBI database |
|---|---|---|---|
| Wave 1 | 1 | - | - |
| Wave 2 | 39 | 7 | 23 |
| Wave 3 | 0 | - | - |
| Wave 4 | 20 | 3 | 12 |
| Wave 5 | 7 | 2 | 2 |
| Wave 6 | 5 | 0 | 1 |
| Wave 7 | 23 | 0 | 2 |
| Wave 8 | 1 | - | - |
